# Cingulin unfolds ZO-1 and organizes myosin-2B and γ-actin to mechanoregulate apical and tight junction membranes

**DOI:** 10.1101/2020.05.14.095364

**Authors:** Ekaterina Vasileva, Florian Rouaud, Domenica Spadaro, Wenmao Huang, Adai Colom, Arielle Flinois, Jimit Shah, Vera Dugina, Christine Chaponnier, Sophie Sluysmans, Isabelle Méan, Lionel Jond, Aurélien Roux, Jie Yan, Sandra Citi

## Abstract

How junctional proteins regulate the mechanics of the plasma membrane and how actin and myosin isoforms are selectively localized at epithelial cell-cell junctions is poorly understood. Here we show by atomic force indentation microscopy, immunofluorescence analysis and FLIM membrane tension imaging that the tight junction (TJ) protein cingulin maintains apical surface stiffness and TJ membrane tortuosity and down-regulates apico-lateral membrane tension in MDCK cells. KO of cingulin in MDCK, mCCD and Eph4 cells results in a decrease in the juxta-membrane accumulation of labeling for cytoplasmic myosin-2B (NM2B), γ-actin, phalloidin and ARHGEF18, but no detectable effect on myosin-2A (NM2A) and β-actin. Loss of paracingulin leads to weaker mechanical phenotypes in MDCK cells, correlating with no detectable effect on the junctional accumulation of myosins and actins. Cingulin and paracingulin form biomolecular condensates, bind to the ZU5 domain of ZO-1, and are recruited as clients into ZO-1 condensates in a ZU5-dependent manner. Cingulin binding to ZO-1 promotes the unfolding of ZO-1, as determined by interaction with DbpA in cells lacking ZO-2 and in vitro. Cingulin promotes the accumulation of a pool of ZO-1 at the TJ and is required in a ZU5-dependent manner for the recruitment of phalloidin-labelled actin filaments into ZO-1 condensates, suggesting that ZU5-cingulin interaction promotes ZO-1 interaction with actin filaments. Our results indicate that cingulin tethers the juxta-membrane and apical branched γ-actin-NM2B network to TJ to modulate ZO-1 conformation and the TJ assembly of a pool of ZO-1 and fine-tune the distribution of forces to apical and TJ membranes.

## INTRODUCTION

The cytoskeleton orchestrates cell shape, motility, internal architecture and mechanical properties, and is involved in most physiological and pathological cellular processes. In epithelial and endothelial tissues, the cytoskeleton is also crucial for the organization and physiology of cell-cell junctions, including tight junctions (TJ), which provide a semipermeable seal for solute diffusion across the paracellular pathway [1, 2], and adherens junctions (AJ), which establish and maintain cell-cell adhesion [3]. Both TJ and AJ are associated with actin and myosin filaments and microtubules, which regulate their dynamic assembly, disassembly and functions [4-7]. Force generated by extracellular and intracellular cues is transduced by junctional mechano-sensing proteins, which modify their conformations and interactions to regulate adhesion strength and cell behavior [8, 9]. TJ and AJ proteins contribute to the organization and function of the cytoskeleton, both through direct interactions with cytoskeletal proteins and through the recruitment of specific activators and inactivators of Rho family GTPases [2, 10].

Most of the mechanical properties of epithelial cells depend on the actin- and myosin-containing cell cortex: myosin-2 ATPase activity and unbranched F-actin polymerization increase cortex tension, while branched actin networks decrease it [11]. Cytoplasmic isoforms of myosin have different roles and localizations at epithelial junctions. Nonmuscle myosin-2B (NM2B) associates with the branched actin meshwork proximal to the plasma membrane, whereas nonmuscle myosin-2A (NM2A), which provides mechanical tugging force, sits on distant peri-junctional actin bundles parallel to the junction [12]. Down-regulation of cytoplasmic β-actin, which is detected laterally and junctionally in epithelial cells, inhibits AJ biogenesis, whereas down-regulation of γ-actin, which is localized apically and junctionally, impairs TJ, but not AJ [13, 14]. However, nothing is known about the mechanisms that direct the selective spatial organization of cytoplasmic actin and myosin isoforms at apical junctions.

The TJ cytoplasmic scaffolding proteins ZO-1 and ZO-2 are critically important not only to link the actomyosin cytoskeleton to TJ transmembrane proteins, but also to control its apical/junctional organization [15-19]. Depletion of ZO-1 results in increased junctional contractility coupled to decreased NM2B integration into junctions [20-22], loss of tortuosity of the TJ membrane [20, 23], increased apical stiffness [24], and altered organization of apical actin filaments [18, 19]. However, the mechanisms through which ZO proteins organize the actomyosin cytoskeleton and mediate its effects on TJ and apical membranes are not well understood. For example, the TJ barrier phenotype in cells lacking ZO-1 can be rescued by constructs lacking the actin-binding region (ABR) within the C-terminal half of ZO-1 [20], suggesting that indirect interactions with actin- and myosin-binding proteins mediate ZO-1 cross-talk with actomyosin. Actomyosin tension and dimerization control the conformation of ZO-1, which can be either stretched (unfolded) or folded (auto-inhibited) [25]. The folded conformation was proposed to result from a mechanosensitive intramolecular interaction between the C-terminal ZU5 (Cter) domain of ZO-1 and the PDZ3-SH3-GUK (PSG) region [25]. Unfolding and multimerization of ZO proteins are required for their liquid-liquid phase separation to drive TJ formation, regulated by phosphorylation and multivalent interactions [26]. Although the N-terminal half of ZO-1 can by itself undergo multimerization and phase separation [26], the C-terminal half of ZO-1 is required to confer mechano-sensitivity to junctions [27], to provide fluidity to ZO-1 condensates [26], and to allow MLCK-dependent regulation of the dynamic behavior of ZO-1 [28]. Together, these observations suggest that ZO-1 interactions mediated by its C-terminal region are critical for ZO-1 mechano-chemical signaling.

Among several actomyosin-associated proteins that bind to ZO-1 [5], cingulin (CGN) [29] and paracingulin (JACOP/CGNL1) [30, 31] interact with actin and myosin [32-35], and with GEFs and GAPs for Rho family GTPases, such as GEF-H1, ARHGEF18 and MgcRacGAP [22, 36-39]. Cingulin is recruited to TJ by ZO-1 [40, 41], whereas paracingulin is recruited to TJ and the *zonula adhaerens* (ZA) through multiple interactions, including with ZO-1 and PLEKHA7 [30, 42]. Cingulin and paracingulin interact with ZO-1 through a conserved ZO-1 Interaction Motif (ZIM) at their N-terminus [32, 40, 42]. Although the sequences of ZO-1 that interact with either cingulin or paracingulin are not known, yeast-2-hybrid screen [43] and BioID experiments [44] indicate that they are within the C-terminal region of ZO-1. Nothing is known about the role of cingulin and paracingulin in the regulation of the mechanical properties of the apical and junctional membranes, the organization of apical actomyosin filaments, and the conformation and junctional assembly of ZO-1. Here we address these questions, by using CRISPR-KO epithelial cell lines and biophysical, biochemical and cellular approaches. We show that cingulin regulates apical surface stiffness and apicolateral membrane tension, junctional accumulation of NM2B and γ-actin, and ZO-1 conformation, TJ assembly and interaction with actin.

## RESULTS

### Knock-out of cingulin decreases apical surface stiffness and increases junctional membrane tension

To ask if cingulin and paracingulin regulate the mechanical properties of epithelial cells, we generated clonal lines of epithelial cells KO for either cingulin, paracingulin or both (Figure S1). Immunoblotting, immunofluorescence and genomic sequence analysis validated the KO lines (Figure S1). In MDCK cells, the single and double CGN/CGNL1 KO clonal lines showed proliferation curves and expression levels of other junctional proteins similar to wild-type (WT), except for an increase in paracingulin levels in CGN-KO cells (Figure S1).

Atomic force indentation microscopy was used to measure the stiffness of the apical surface in MDCK clonal lines (Figure 1A). Force-indentation curves of the MDCK cells were fitted by Hertz model [45] (Figure 1B) to obtain the Young’s modulus (Figure 1C). The Young’s modulus of CGN-KO and CGN/CGNL1 double-KO MDCK cells was less than half the value of WT cells (e.g. 0.0015 MPa and 0.0014 MPa, compared to 0.0037 MPa) (Figure 1C and Table S1), indicating a significant loss of stiffness of CGN-KO and double-KO cells. The decrease in Young’s modulus for CGNL1-KO MDCK cells was smaller (0.0021 MPa), indicating that paracingulin is less important in regulating apical stiffness, and that cingulin is epistatic to CGNL1.

**Figure 1.**
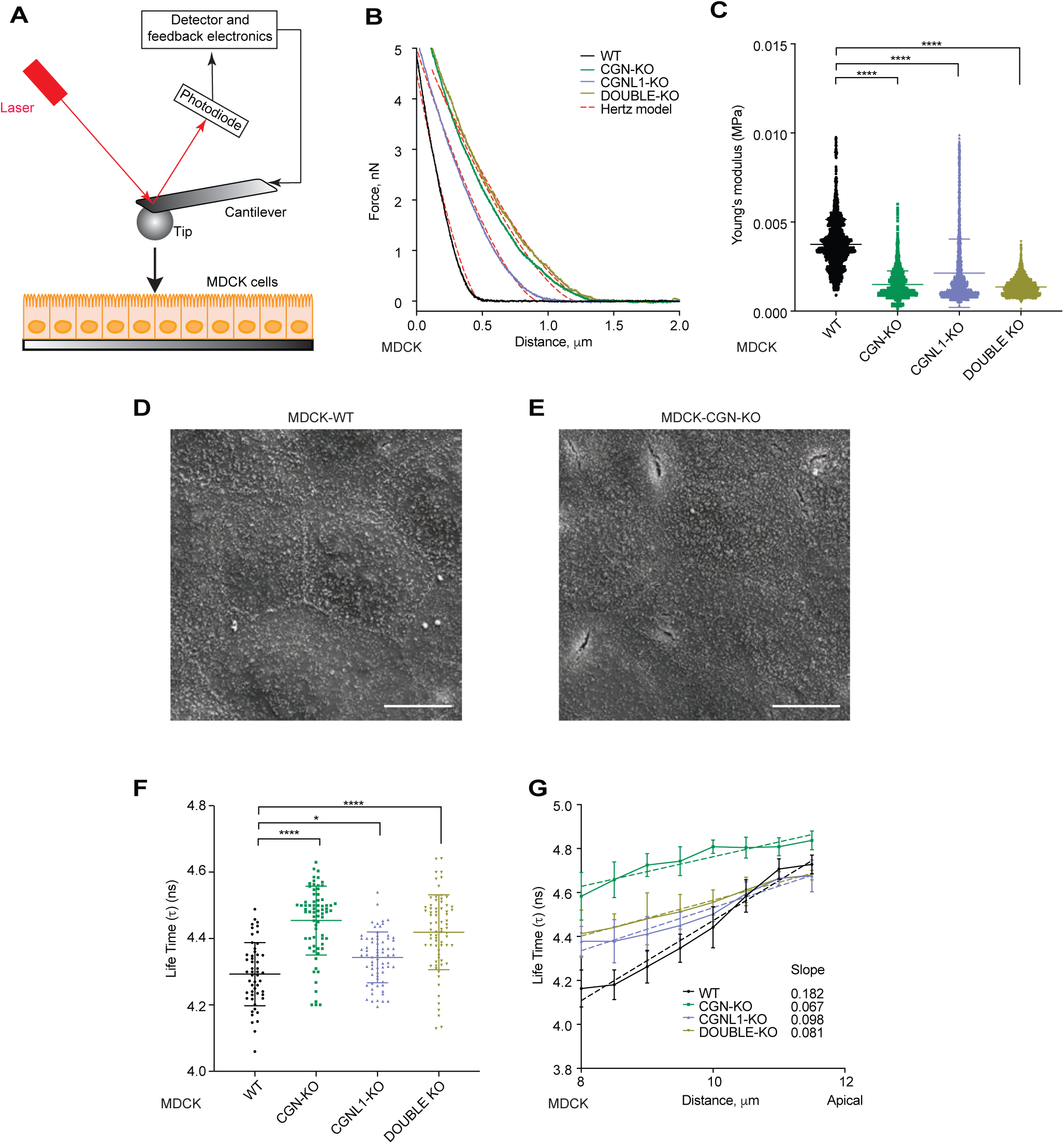
Cingulin KO results in decreased apical membrane stiffness and increased apicolateral membrane tension. (A) Schematic diagram of experimental setup. (B) Representative force-indentation curves of WT, CGN-KO, CGNL1-KO and CGN/CGNL1-double KO MDCK cell lines fitted with Hertz (Spherical) model. (C) Averaged stiffness (Young’s modulus) of WT, CGN-KO, CGNL1-KO and CGN/CGNL1-double KO MDCK cell lines. (D-E) Representative SEM images of MDCK WT (D) and CGN-KO (E) cell lines. Scale bars, 10 μm. (F) Average lifetimes of the FliptR probe for WT, CGN-KO, CGNL1-KO and CGN/CGNL1-double KO MDCK cell lines. (G) FliptR lifetime changes along the apicolateral junctional membrane of WT, CGN-KO, CGNL1-KO and CGN/CGNL1-double KO MDCK cell lines. Slopes of fitted linear regression curves (dashed lines) are indicated. Bars (C, F, G) show mean±SD. Related Figure S1 shows phenotypic characterization of MDCK KO cells.

The shape of the apical surface of CGN-KO cells was examined by scanning electron microscopy of confluent MDCK monolayers. WT cells displayed a slightly extruding, dome-shaped apical surface (Figure 1D), whereas the apical surface of CGN-KO cells appeared flatter (Figure 1E).

Next, we used the FliptR membrane probe [46] to measure membrane tension along the apico-lateral region of the plasma membrane of MDCK cells. Probe lifetime, which correlates with membrane tension [46], was strongly increased in CGN-KO and double-KO MDCK cells, and slightly, but still significantly increased in CGNL1-KO cells (Figure 1F). By analyzing lifetime emission of the FliptR probe as a function of the distance from the apical surface, WT cells showed a sharp gradient of tension along 4 μm, from the basal to the apical surface (slope 0.182, Figure 1G), whereas the gradient for cingulin-KO cells was less sharp (slope 0.067, Figure 1G), indicating increased tension throughout this region. The slopes for CGNL1-KO and double KO cells were 0.098 and 0.081, respectively (Figure 1G) suggesting a less important role of paracingulin in the maintenance of a sharp membrane tension gradient.

Together, these observations indicate that cingulin, and to a lesser extent paracingulin, is required to maintain the stiffness of the apical surface membrane and down-regulate tension at the apico-lateral, junctional membrane.

### Cingulin maintains the tortuosity of the TJ membrane and is required for the correct organization of nonmuscle myosin-2B (NM2B) and γ-actin at junctions

To understand how cingulin and paracingulin regulate the mechanical properties of the apicolateral plasma membrane, we examined the shape of the TJ membrane in MDCK cells and the spatial organization of cytoplasmic myosin and actin isoforms in different cell types KO for one of both proteins. In polarized WT MDCK cells labeling for cingulin and occludin was wavy, whereas β−catenin labeling, corresponding to the *zonula adhaerens* (ZA) and the AJ, was straight (Figure 2A, and Figure 2B-WT), suggesting that the membranes of TJ and ZA/AJ are subjected to different tensile forces. To quantify TJ membrane tortuosity, we used the zigzag index [23]. KO of either cingulin or both cingulin and paracingulin resulted in straight TJ membranes (Figure 2B) and a large drop of the zig-zag index (Figure 2C), whereas KO of paracingulin resulted in a considerably smaller reduction in the zig-zag index (Figure 2B-C). The decreased TJ membrane tortuosity of CGN-KO MDCK cells was rescued by re-expression of either canine, mouse or human cingulin (Figure 2D-E, Figure S2A, S2C), but not by overexpression of ZO-1 (Figure S2A, S2C). Furthermore, tortuosity of the TJ membrane in the background of WT cells was increased by overexpression either of cingulin or ZO-1 (Figure S2A-B), the latter being associated with increased junctional cingulin (Figure S2D).

**Figure 2.**
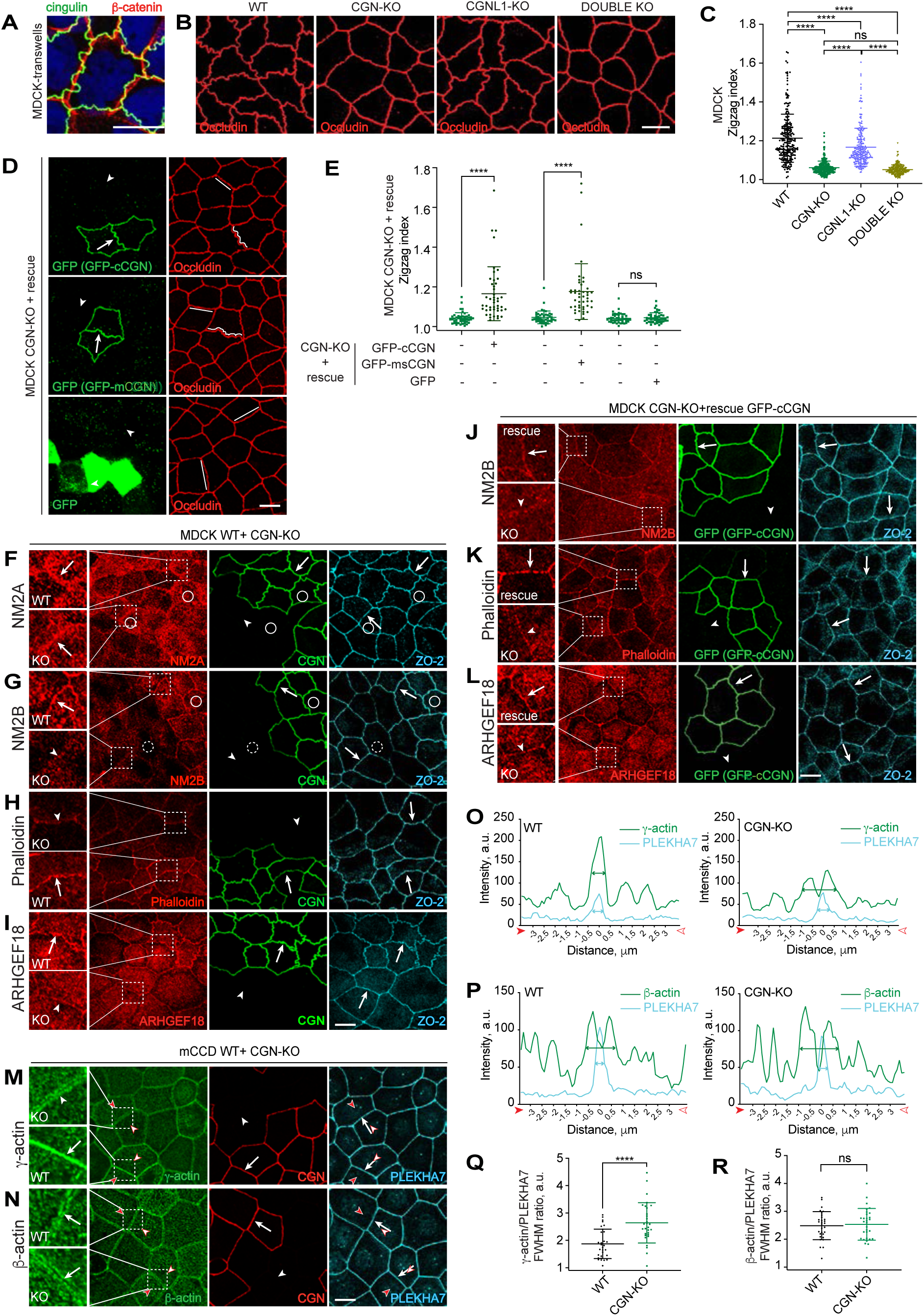
Cingulin is required to maintain membrane tortuosity, and normal accumulation of myosin-2B, actin and ARHGEF18 at TJ. (A-B) Immunofluorescent localization of β-catenin and cingulin in WT cells (A) and occludin in WT and KO lines (B) using MDCK cells grown on Transwell filters. (C) Quantifications of membrane tortuosity (zigzag index) in WT, CGN-KO, CGNL1-KO and CGN/CGNL1-double KO MDCK cell lines. (D-E) Immunofluorescent localization of occludin and GFP-c/mCGN/GFP (D) and quantifications of membrane tortuosity (zigzag index) (E) in CGN-KO MDCK cells rescued with either GFP-canisCGN (cCGN), or GFP-mouseCGN (mCGN) or GFP. Arrows and arrowheads indicate junctional localization and lack of junctional localization, respectively. White lines outline shape of junctions. (F-I) Immunofluorescent localization of NM2A (F), NM2B (G), F-actin (TRITC-phalloidin) (H) and ARHGEF18 (I) in mixed WT+CGN-KO MDCK cultures. Arrows and arrowheads indicating normal versus reduced/undetectable labeling are placed in magnified areas for the red channel and in regular magnification areas in CGN and ZO-1 channels. Continuous and dotted circles indicate normal and reduced, respectively, apical cortical labeling for NM2A and NM2B. (J-L) Immunofluorescent localization of NM2B (J), F-actin (TRITC-phalloidin) (K) and ARHGEF18 (L) in CGN-KO cells rescued with full length canis cingulin. (M-R) Immunofluorescent localization (M-N) and line-scans of immunofluorescence signal across junctions (O-P) of γ-actin (M, O) and β-actin (N, P) in junctions from either WT or KO cells within mixed WT-CGN-KO mCCD cultures. Arrows and arrowheads in magnified insets show normal and altered junctional labeling, respectively. (Q-R) show the ratio between the full width at half maximum signal intensity (FWHM) for γ-actin (Q) and β-actin (R), ratioed to PLEKHA7 signal (cyan) (N=30 junctions). Data in (C, E, Q, R) are represented as mean±SD. Scale bars= 10 μm. Related Figure S1 shows phenotypic characterization of MDCK and mCCD KO cells. Related Figure S2 shows analysis of zigzag index, NM2A, NM2B, phalloidin, ARHGEF18, γ-actin and β-actin in different cell types and experimental conditions. Related Figure S6D shows schematic diagram of interaction of cingulin with ZO-1, γ-actin filaments and NM2B.

Next, we examined the distribution of myosins and actins in mixed cultures of WT and KO cells, using antibodies against cytoplasmic nonmuscle myosin-2A (NM2A), myosin-2B (NM2B), γ-actin and β-actin isoforms, and fluorescently labeled phalloidin. In WT MDCK cells NM2A and NM2B showed a junctional localization (arrows in Figure 2F-G, insets), and a diffuse apical distribution (circles in Figure 2F-G), this latter likely corresponding to the terminal web and apical cortex. In CGN-KO MDCK cells, labeling for NM2A was similar to WT (arrow, Figure 2F), whereas in CGN-KO cells there was a decrease in both junctional and apical NM2B labeling (arrowhead and dotted circle, Figure 2G), as well as a decrease in junctional labeling for phalloidin (arrowhead, Figure 2H). A similar decrease in NM2B and phalloidin junctional labeling and no detectable change in NM2A labeling was observed using CGN-KO+WT mixed cultures of Eph4 cells (Figure S2G) and mCCD cells (Figure S2K). However, the TJ membrane loss-of-tortuosity phenotype could not be detected in Eph4 and mCCD cells, since both WT and CGN-KO cells had straight TJ membranes (Figure S2G-L). The junctional accumulation of ARHGEF18 (p114RhoGEF), which regulates Rho signaling and actomyosin organization at junctions [39], was also decreased in CGN-KO MDCK cells (arrowhead, Figure 2I). NM2B, phalloidin and ARHGEF18 labeling were rescued in CGN-KO cells by re-expression of GFP-tagged cingulin (arrows, Figure 2J-L), whereas GFP alone had no effect (arrowheads, Figure S2F). In contrast, in mixed cultures of WT and CGNL1-KO WT cells there were no clearly detectable differences in the accumulation of NM2A, NM2B, phalloidin and ARHGEF18 at junctions (arrows, Figure S2E). Next, we investigated the distribution of cytoplasmic β-actin and γ-actin isoforms, using specific antibodies [13]. Since the labeling of MDCK cells with these antibodies was less clear, we used mCCD and Eph4 cells. Strikingly, junctional labeling of γ-actin, which was intense in the juxtamembrane region of WT cells, was reduced in intensity and shifted farther away from the membrane in CGN-KO mCCD cells (arrowhead, Figure 2M inset). In contrast, the distribution of labeling for β-actin, which was farther away from the membrane than γ-actin, was similar in WT and CGN-KO mCCD cells (arrows, Figure 2N). Quantification of line scans of labeling across the junction and colocalization with PLEKHA7 confirmed that KO of cingulin resulted in a shift of γ-actin labeling farther from the membrane, with respect to WT cells (Figure 2O-Q), and no detectable change for β-actin (Figure 2P-R). Similar results were obtained in Eph4 cells, where KO of cingulin resulted in decreased junctional labeling for NM2B, phalloidin and γ-actin, but no detectable change in NM2A and β-actin labeling (arrows and arrowheads in Figure S2G). Similar to MDCK cells, KO of paracingulin in Eph4 and mCCD cells did not result in clearly detectable differences in labeling for myosins, phalloidin and actins (Figure S2H, S2L). The decreased junctional localization of NM2B and F-actin in CGN-KO Eph4 cells was rescued by re-expression of cingulin (Figure S2I-J), and of chimeric constructs comprising the head domain of CGNL1 and the rod+tail domain of CGN (Figure S2J) [30, 32], indicating that the rod+tail region of cingulin is mechanistically involved. In summary, cingulin plays a key role in maintaining membrane tortuosity and organizing NM2B and γ-actin at TJ of epithelial cells, whereas paracingulin has a minor role in maintaining membrane tortuosity in MDCK cells and does not induce clearly detectable changes in the organization of actin and myosin filaments in this experimental model.

### Cingulin binding to the ZU5 domain of ZO-1 unfolds ZO-1

To investigate the functional consequences of the cingulin-dependent organization of NM2B and γ-actin at TJ, we focused on ZO-1, the major protein implicated in TJ-actin interactions, and the protein that recruits cingulin to TJ. Specifically, since ZO-1 monomers undergo tension-dependent stretching (unfolding) and folding [25], we hypothesized that by promoting the accumulation of NM2B near TJ, cingulin might facilitate the transduction of force onto ZO-1 monomers, and thus promote their unfolding. Alternatively, cingulin may affect ZO-1 conformation by binding to a region involved in intramolecular interactions, such as the C-terminal ZU5 domain (Cter, [25]).

To test these hypotheses, we first mapped the region of ZO-1 that binds to cingulin. A N-terminal fragment of cingulin that contains the ZO-1-Interaction-Motif (ZIM) (CGN(1-70)) [40] was used as a bait to pull down GFP-tagged fragments of the C-terminus of ZO-1 (residues 888-1748)(Figure S3A). Only ZO-1 preys that contained the ZU5 domain (ZU5=Cter, residues 1619-1748 [25]) interacted with GST-CGN(1-70) (Figure 3A), and the binding of the ZIM to the isolated ZU5 fragment (1619-1748) was >4-fold stronger than to other fragments (Figure S3B). Deleting the ZIM-containing first 70 residues of cingulin (CGN-Δ1-70) essentially abolished the interaction between bacterially expressed ZU5 and full length cingulin (Figure 3B). Fragmenting the ZU5 domain strongly decreased the interaction with the CGN-(1-70) bait (Figure S3 C-F), indicating that an intact ZU5 domain is required for high affinity binding of ZO-1 to cingulin. Full-length paracingulin also interacted, albeit weaker than cingulin, with the ZU5 bait, and deleting its first 80 residues abolished the interaction (Figure 3C). Larger ZIM-containing fragments of paracingulin also showed weaker binding to ZO-1, with respect to cingulin (Figure S3G). Junctional cingulin labeling was undetectable in ZO-1-KO cells (Figure S3H) (see also [41]), whereas paracingulin labeling was only reduced, but not abolished in ZO-1-KO Eph4 cells (Figure S3I-J). The loss of cingulin labeling in ZO-1-KO cells was rescued by expressing full-length ZO-1, but not ZO-1 lacking the ZU5 domain (ZO-1-ΔZU5) (Figure 3D). As an alternative approach to test ZO-1 binding to cingulin and paracingulin, we examined their recruitment in ZO-1 condensates obtained by exogenous overexpression of ZO-1 [26]. Both cingulin and paracingulin were recruited into ZO-1 condensates in a ZU5-dependent manner (Figure 3E-F). Together, these results demonstrate that the ZU5 domain of ZO-1 binds to cingulin and paracingulin and indicate that only a fraction of paracingulin is recruited to TJ by ZO-1, consistent with previous observations [30, 42].

**Figure 3.**
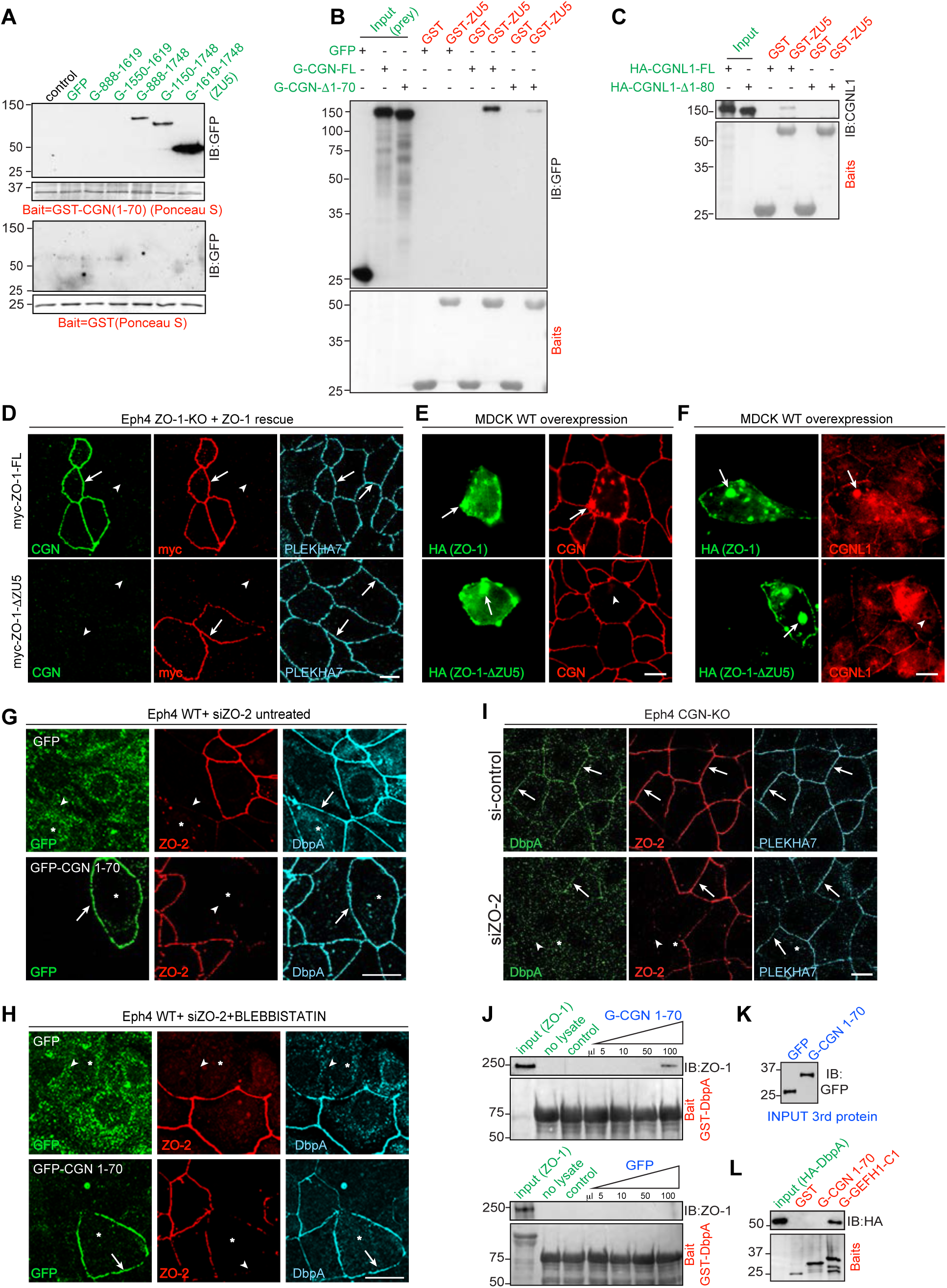
Cingulin binding to the ZU5 domain unfolds ZO-1. (A-B) Interaction between the GST-CGN(1-70) bait (red) and GFP-tagged fragments of ZO-1 (input preys, green) (A), and between the GST-ZU5 bait and GFP-tagged cingulin preys (B), as shown by IB with anti-GFP. Numbers indicate migration of pre-stained size markers. PonceauS-stained baits are shown below the IB. GST alone is used as a negative control in all GST pulldown experiments. (C) Interaction between the GST-ZU5 bait and HA-tagged paracingulin full length and mutated (CGNL1) preys. (D) Immunofluorescent localization of endogenous CGN (CGN, green), myc-tagged ZO-1 (red) and PLEKHA7 (reference junctional marker, cyan) in ZO-1-KO Eph4 cells rescued either with FL-ZO-1 (top) or with a C-terminal truncation of ZO-1 lacking the ZU5 domain (bottom). Arrows and arrowheads indicate normal and reduced/undetected staining, respectively. (E-F) Recruitment of CGN (E) and CGNL1 (F) in condensates of full-length myc-ZO-1-HA (top), but not in condensates of ZO-1 lacking the ZU5 domain (ZO-1-ΔZU5) (bottom). (G-I) CGN(1-70) promotes ZO-1 unfolding in cells. Junctional recruitment of DbpA in WT Eph4 cells depleted of ZO-2 and overexpressing ether GFP or GFP-CGN(1-70) (G-H) either untreated (G) or treated with blebbistatin (H), and in CGN-KO Eph4 cells treated with either si-control or si-ZO2 (I). Asterisks indicate ZO-2-depleted cells. Arrows and arrowheads indicate normal and reduced/undetected junctional staining, respectively. (J-L). Cingulin (1-70) promotes ZO-1 unfolding in vitro. Immunoblot analysis (J) of full-length ZO-1 prey (purified from baculovirus-infected insect cells) in GST pulldowns using GST-DbpA as a bait and increasing amounts of either GFP-CGN(1-70) or GFP as third protein. (K) shows normalization of third protein. (L) Immunoblot analysis of HA-DbpA prey in GST pulldowns using GST-CGN(1-70) and GST-GEF-H1 as baits [47]. Scale bars, 10 μm. Related Figure S1 shows phenotypic characterization of Eph4 KO cells. Related Figure S3 shows binding of cingulin and paracingulin fragments to ZU5, localization of CGN and CGNL1 in ZO-1-KO Eph4 cells, and measurement of the K_d_ of interaction between GST-CGN(1-70) and ZU5. Related Figure S6 shows schematic diagram of how cingulin binds to and unfolds the ZU5 domain of ZO-1, thus connecting it to actin filaments.

Second, we asked whether cingulin binding to the ZU5 domain promotes ZO-1 unfolding. We proposed that when ZO-1 is in the folded conformation, the ZU5 domain binds to the PDZ3-SH3-GUK region of ZO-1 (ZPSG1) [25]. Thus, we wondered whether cingulin binding induces ZO-1 unfolding by competing with ZU5-dependent intramolecular interactions. To test this, we overexpressed the (1-70) fragment of cingulin, which contains the ZO-1-binding ZIM motif, in Eph4 cells, and used the junctional localization of DbpA as a readout for the ZO-1 stretched conformation [25]. ZO-1 was sufficient to retain DbpA at junctions in cells depleted of ZO-2 (arrow Figure 3G, upper panels), but DbpA junctional localization was reduced or undetectable in cells depleted of ZO-2 and treated with the myosin inhibitor blebbistatin (arrowhead, Figure 3H, upper panel), as shown previously [25]. GFP-CGN(1-70) was recruited to junctions (Figure 3G, bottom panel), and its overexpression in ZO-2-depleted cells treated with blebbistatin rescued the junctional localization of DbpA (arrow in Figure 3H, bottom panel), indicating ZO-1 unfolding [25]. To confirm the role of cingulin in unfolding ZO-1, we used CGN-KO Eph4 cells. Depletion of ZO-2 in these cells resulted in loss of DbpA at junctions (arrowhead, Figure 3I, bottom panel), indicating that loss of cingulin phenocopies treatment with blebbistatin of ZO-2-depleted cells [25]. Furthermore, although GST-DbpA failed to interact with full-length ZO-1 in vitro [25, 47], addition of increasing amounts of GFP-CGN(1-70) resulted in detectable ZO-1 in GST-DbpA pulldowns (Figure 3J, normalization of preys in Figure 3K), indicating ZO-1 unfolding. This interaction, and the rescue of junctional DbpA by CGN(1-70) (Figure 3H) was not an artefact due to binding of CGN(1-70) to DbpA, since this construct did not interact with DbpA, which could still bind to its known interactor GEF-H1 [48], used as a positive control (Figure 3L). The dissociation equilibrium constant (K_d_) for the interaction of the CGN(1-70) with ZU5 was 40.4 nM (Figure S3K-M), lower that the calculated K_d_ for the ZPSG-ZU5 interaction (66 nM, [25]). Collectively, these experiments indicate that high affinity binding of the ZIM-containing region of cingulin to the ZU5 domain of ZO-1 is required for cingulin recruitment to junctions and promotes the unfolding of ZO-1.

### Cingulin promotes the accumulation of ZO-1 at tight junctions

Since ZO-1 unfolding promotes phase separation and TJ assembly of ZO-1 [26], we hypothesized that cingulin, which is recruited to TJ by ZO-1, could in turn enhance the assembly of ZO-1 at TJ, in a positive feedback loop. To test this, we compared ZO-1 immunofluorescent labeling in confluent monolayers of WT cells, and cells KO for cingulin. Using either the ZA protein PLEKHA7 or the TJ protein occludin as a reference junctional marker, ZO-1 labeling at apical junctions was reduced in CGN-KO cells, when compared to WT, in mixed cultures of WT+CGN-KO Eph4 cells (Figure 4A, Figure S4A-B) and MDCK cells (Figure 4B). In contrast, ZO-1 labeling was similar in mixed cultures of WT and CGNL1-KO Eph4 (Figure 4C) and MDCK cells (Figure 4D). The same results were obtained when examining the effect of KO of either CGN or CGNL1 in mCCD cells (Figure S4C-D, G-H), and when labeling ZO-1 with different antibodies (Figure S4A-H), indicating that this phenotype is not affected by cell type or epitope availability differences. Next, to confirm that the reduction in ZO-1 labeling in CGN-KO cells was specifically due to the loss of cingulin, and not to clone-dependent variations, we rescued the localization of ZO-1 through expression of exogenous cingulin constructs. Full-length, myc-tagged cingulin rescued ZO-1 junctional accumulation (arrows in Figure 4E, CGN FL-myc, and quantification in Figure 4G), whereas a mutant of cingulin lacking the first 70 amino acid residues of cingulin, which contain the ZO-1-Interaction-Motif (ZIM) [40] did not localize at junctions, and did not rescue ZO-1 junctional labeling (arrowheads, Figure 4E, CGN-Δ1-70-myc, quantification in Figure 4G). Conversely, highly overexpressed full-length cingulin, but not cingulin lacking the first 70 residues, enhanced the junctional accumulation of ZO-1 in otherwise WT cells (Figure 4F), even though this increase was not statistically significant (Figure 4G), suggesting that in WT cells ZO-1 levels are already near saturation. By immunoblotting analysis, the KO of either CGN or CGNL1 did not affect the total levels of ZO-1 protein and its α(+) versus α(-) isoforms (Figure S1R, S4I-J), indicating that the reduced junctional labeling of ZO-1 is not due protein degradation or to a loss of a specific isoform.

**Figure 4.**
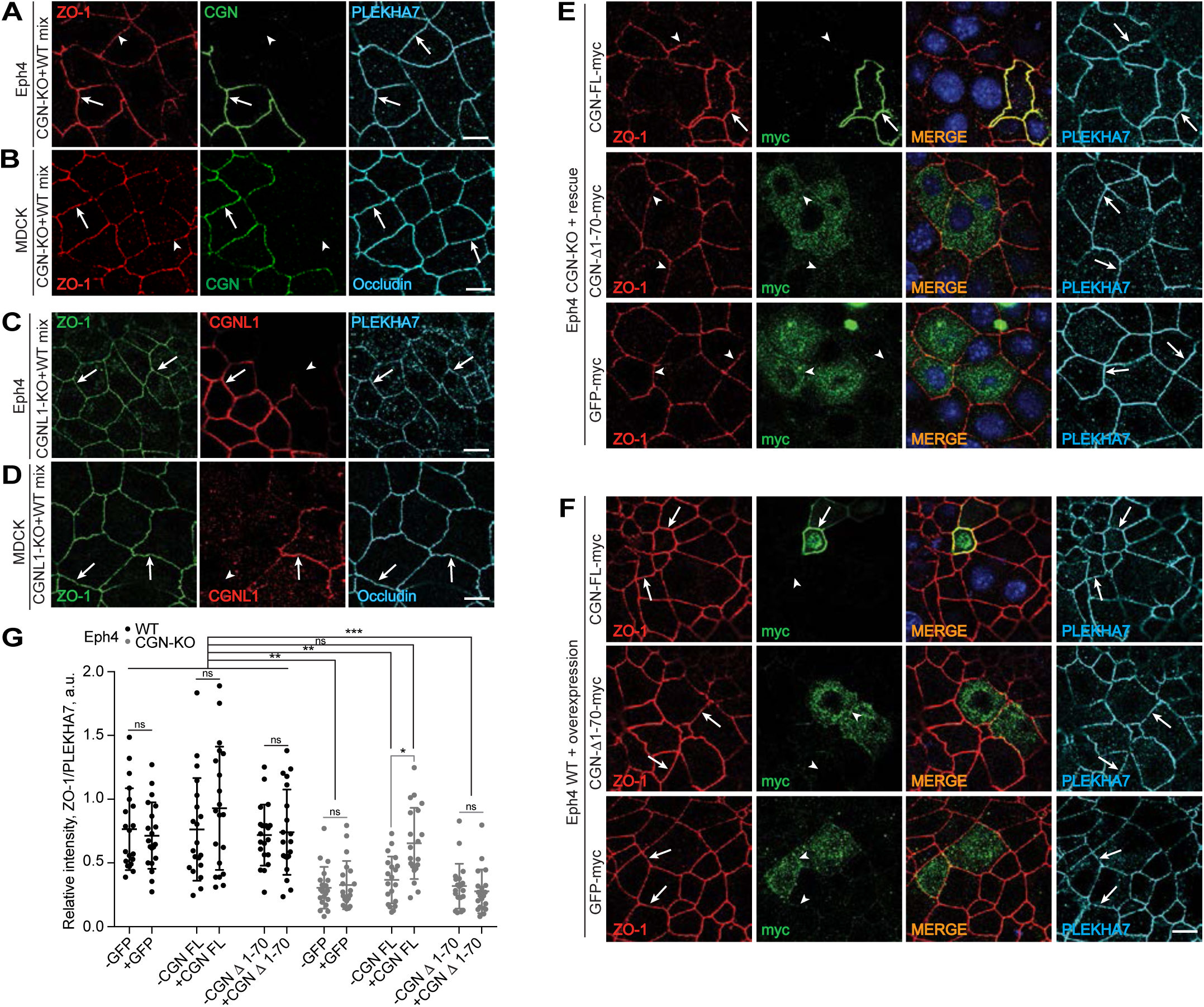
Cingulin promotes the accumulation of a pool of ZO-1 at TJ. (A-D) KO of cingulin (A-B) but not paracingulin (C-D) decreases the junctional accumulation of ZO-1 in Eph4 (A, C) and MDCK (B, D) cells. (E-F) Immunofluorescent localization of ZO-1 (ZO-1), CGN rescue constructs full length (CGN-FL), mutated (CGN-Δ1-70) and GFP myc-tagged constructs (myc) and PLEKHA7 (PLEKHA7) in either CGN-KO (E) or WT (F) cultures of Eph4 cells. Arrows and arrowheads indicate normal and reduced/undetected junctional labeling, respectively. Scale bars, 10 μm. (G) Quantification of ZO-1 junctional immunofluorescence signal in either WT or CGN-KO cells after rescue with either GFP-myc or full-length or mutated CGN constructs (E-F). Data are represented as mean±SD. Related Figure S1 shows phenotypic characterization of KO cells. Related Figure S4 shows IF analysis and quantifications of IF labeling for ZO proteins in additional cell lines, and IB analyses.

We next asked whether other ZO proteins are affected by the loss of cingulin. The junctional labeling for ZO-2 was similar to WT in cells KO for either CGN or CGNL1 (Figure S4K-L), whereas ZO-3 labeling was reduced in CGN-KO cells (Figure S4O-P), but not in CGNL1-KO cells (Figure S4Q-R). ZO-3 labeling was rescued by re-expression of cingulin, but not overexpression of ZO-1 (Figure S4U-V). The protein expression levels of ZO-2 and ZO-3 were not affected by KO of CGN either in MDCK cells (Figure S1R, S4M-N, S4S-T), or mCCD and Eph4 cells (S4M-N, S4S-T). Together, these data indicate that cingulin controls the accumulation of a pool of ZO-1 and ZO-3, but not ZO-2, at TJ.

### ZO-1 condensates recruit NM2B and ARHGEF18 independently of cingulin, and cingulin promotes interaction of ZO-1 with F-actin by interacting with the ZU5 domain

To gain more insight into the mechanism through which cingulin promotes the junctional accumulation of ZO-1, we overexpressed GFP-tagged forms of cingulin, paracingulin and ZO-1 in MDCK cells and analyzed the labeling of clients/interactors into phase-separated ZO-1 condensates (Figure 5 and Figure S5) [26]. ZO-1 undergoes liquid-liquid phase separation [26], and previous observations indicate that either cingulin or paracingulin overexpression in MDCK cells leads to the formation of coalescing dots, with dynamics that suggest phase separation [49]. GFP-tagged cingulin and paracingulin, when overexpressed in cells, induced the formation of cytoplasmic brightly labelled structures, which underwent dynamic fission and fusion, (Figure 5A-A’’ and Figure S5A-A’), very similar to what observed for ZO-1 (Figure 5B-B’’), suggesting the formation of phase-separated condensates. We found that while ZO-1 condensates recruited both cingulin and paracingulin (Figure 3E-F) (see also [26]), cingulin and paracingulin condensates did not recruit ZO-1 as a client (Figure S5B). Moreover, NM2B labeling was increased and detected in bright cytoplasmic condensates of either CGN or ZO-1 (arrows, Figure 5C-D). In contrast, in the case of phalloidin and ARHGEF18, labeling was redistributed in the cortical submembrane area of cells that overexpressed CGN (double arrowheads, Figure 5C) and detected in strongly labelled cytoplasmic condensates of ZO-1 (arrows, Figure 5D). Paracingulin condensates also showed redistribution of ARHGEF18 in the submembrane cortex but not in the cytoplasm, similarly to cingulin, but did not show increased or redistributed labeling for either NM2B or phalloidin (Figure S5C). NM2B and ARHGEF18 were brightly labelled in cytoplasmic ZO-1 condensates even when ZO-1 was overexpressed in the background of CGN-KO MDCK cells (Figure 5E), indicating that the association of ZO-1 with NM2B and ARHGEF18 occurs independently of cingulin, at least in the context of condensates. Importantly, the recruitment of phalloidin-labelled F-actin into ZO-1 condensates was significantly decreased in CGN-KO cells, as shown by the lack of brightly labelled cytoplasmic dots (arrowhead, Figure 5E, middle panels), suggesting that cingulin promotes ZO-1 interaction with actin filaments. To test this hypothesis, we compared phalloidin staining in condensates of either full length ZO-1, or of the ZO-1 mutant lacking the ZU5 domain (ZO-1-ΔZU5). Strikingly, while in WT cells condensates made by overexpressing either full-length ZO-1 or ZO-1-ΔZU5 recruited brightly phalloidin-labelled F-actin (arrows, Figure 5F, quantification in Figure 5H), in the background of CGN-KO cells only the ZO-1-ΔZU5 construct, but not full-length ZO-1 could recruit bright phalloidin-labelled F-actin (arrowhead and arrow, Figure 5G, quantification in Figure 5H). This suggests that in full-length ZO-1 the ZU5 domain inhibits ZO-1-F-actin interaction, and that cingulin binding to the ZU5 antagonizes this inhibition.

**Figure 5.**
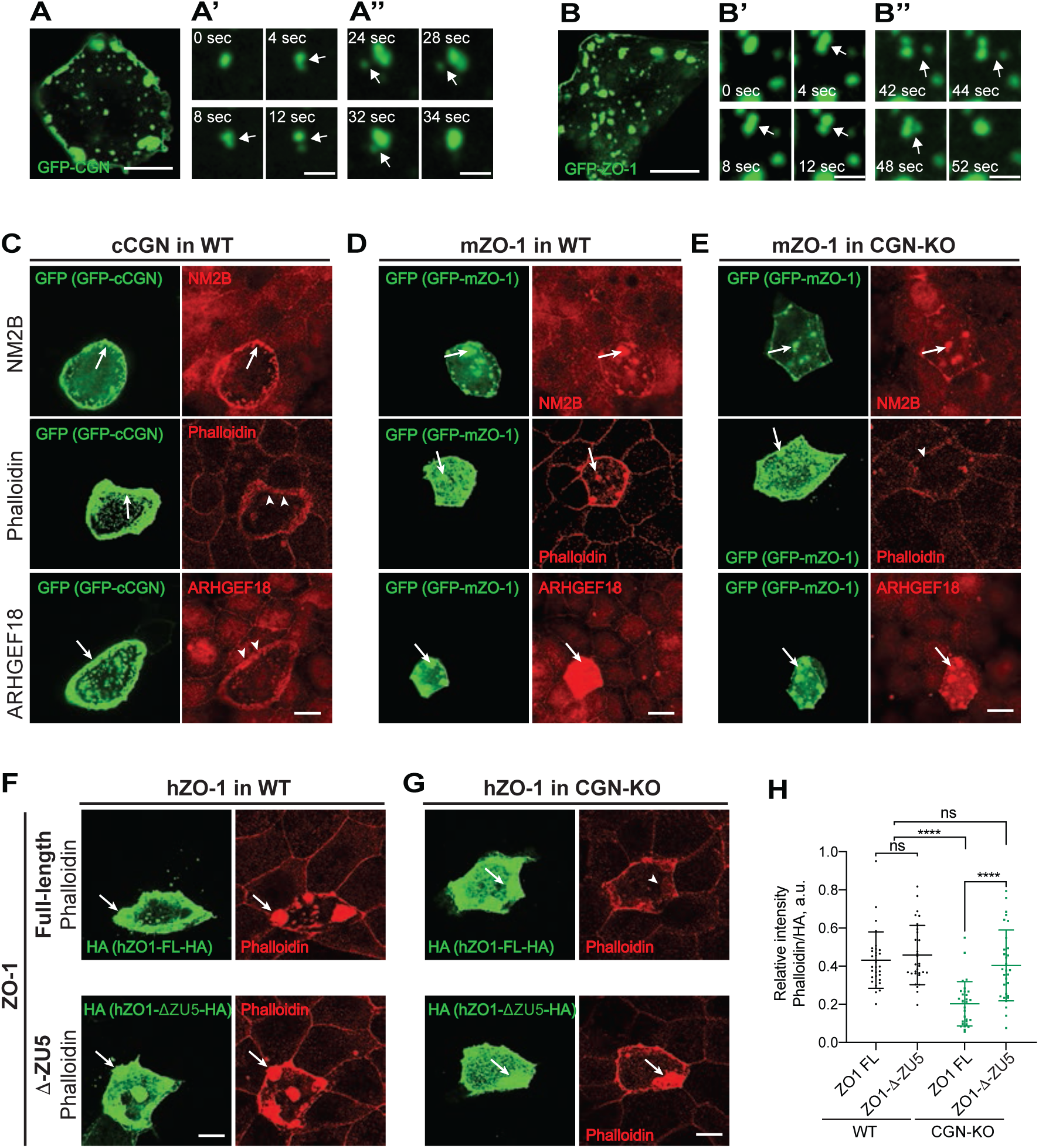
Cingulin is required for recruitment of F-actin but not NM2B to ZO-1 condensates. (A-B) CGN (A) and ZO-1 (B) both form round droplets that fuse into larger structures over time (A’-B’), indicating the formation of condensates. (C-E) Labeling for NM2B (upper panels), F-actin (TRITC-phalloidin) (middle panels) and ARHGEF18 (bottom panels) in cell overexpressing either CGN in MDCK WT (C), or ZO-1 in WT (D) or CGN-KO (E) MDCK cells. (F-G) Recruitment of F-actin (TRITC-phalloidin) in condensates of full-length myc-ZO-1-HA (top) and ZO-1 lacking the ZU5 domain (ZO-1-ΔZU5-HA) (bottom) in WT (F) or CGN-KO (G) MDCK cells. Arrows indicate phase-separated condensates, arrowheads indicate reduced/undetected labeling, double arrowheads indicated redistributed subcortical labeling. (A, B, C-G) Scale bars, 10 μm. (A’, B’) Scale bars, 0.05 μm. (H) Quantification of relative fluorescent intensity of F-actin recruited to ZO-1 condensates in WT and CGN-KO MDCKII (F-G). Data are represented as mean±SD. Related Figure S1 shows phenotypic characterization of MDCK CGN-KO cells. Related Figure S5 shows IF analysis of ZO-1 in CGN and CGNL1 condensates, and NM2B, phalloidin and ARHGEF18 labeling in CGNL1 condensates. Related Figure S6C-D shows schematic diagram of how ZU5 folds back on the ABR in the absence of cingulin, how cingulin binding opens the ZU5 domain, thus allowing the ABR to connect to actin filaments.

## DISCUSSION

The mechanical properties of cells are crucial to respond to environmental cues during development, in adult tissue homeostasis, and in disease, and depend on actin and myosin, and their mode of polymerization [11]. Here we provide evidence that cingulin regulates the mechanical properties of the apical membrane, the junctional organization of γ-actin and NM2B, and the conformation, TJ assembly and F-actin interaction of ZO-1 (Figure 6, Figure S6).

**Figure 6.**
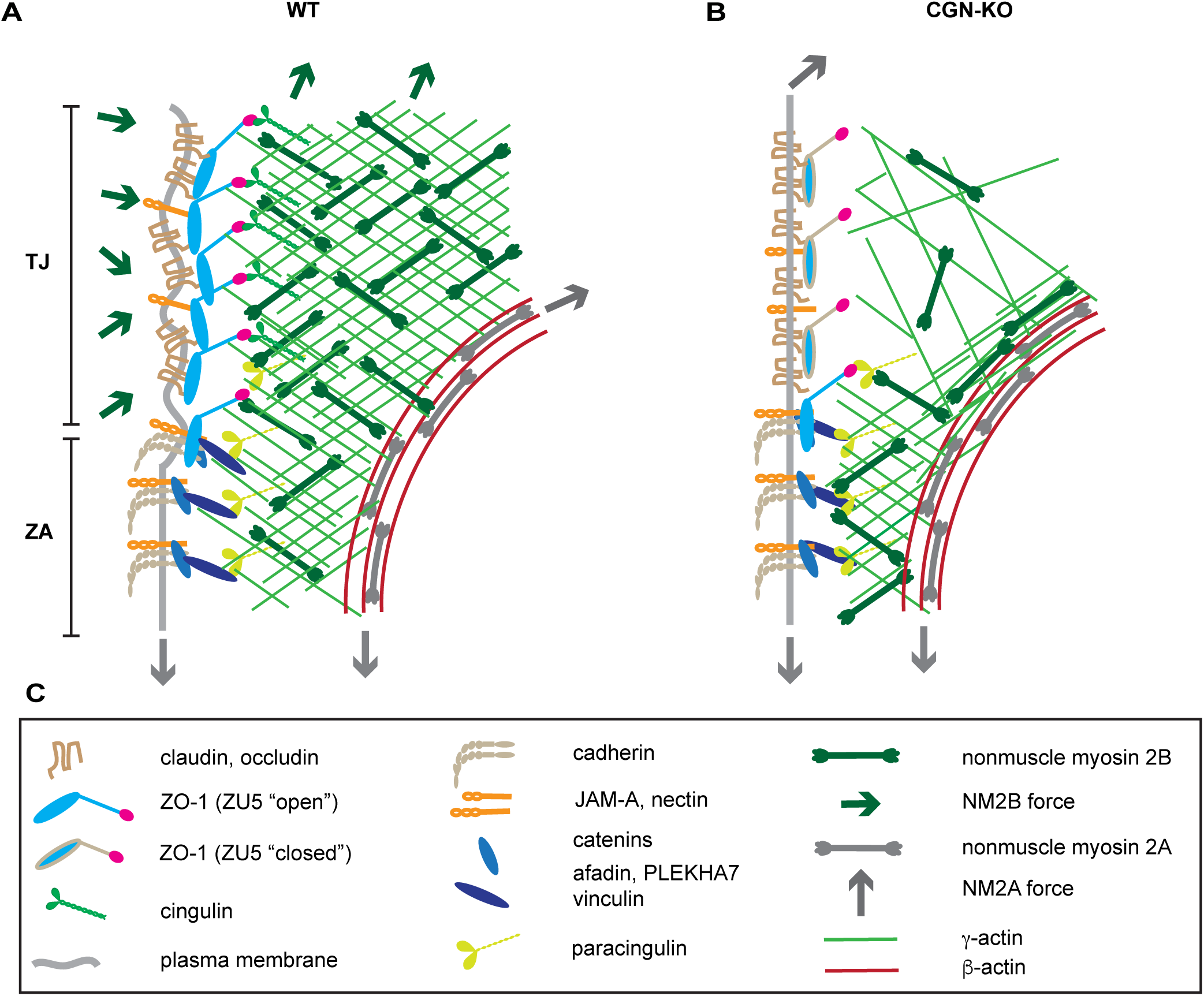
The effects of cingulin KO on membrane shape and cytoskeleton organization. (A-B) Schematic hypothetical organization of cingulin, paracingulin, myosin and actin filaments in the apicolateral junctional region of WT (A) and CGN-KO (B) cells. Selected transmembrane and junctional proteins of tight junctions (TJ) and *zonula adhaerens* (ZA) are shown (see graphical legend). In WT cells γ-actin is tethered to the ZO-1-cingulin complex, and this link is lost in CGN-KO cells, resulting in thinning of the branched γ-actin network and uncoupling from the TJ membrane, straightening of the TJ membrane, and reduced accumulation of ZO-1. (C) Graphical legend. Note that many additional transmembrane and cytoplasmic junctional proteins of TJ, such as ZO-2 and ZO-3, which form heterodimers with ZO-1 have been omitted from the scheme, for the sake of clarity. ZO-1 is represented in “open” and “closed” forms, depending on the conformation and intramolecular interactions of the ZU5 domain, regulated by interaction with cingulin. In cells that contain ZO-2, ZO-1 is unfolded through dimerization. Related Figure S6 shows an additional scheme detailing the folding/unfolding of soluble and membrane-associated pools of ZO-1, its interactions with TJ transmembrane and other proteins, and cingulin-mediated interaction with NM2B and γ-actin.

The observation that straight TJ and reduced junctional accumulation of NM2B [20, 21, 23] occur in CGN-KO cells in the presence of ZO-1 suggests that ZO-1 controls TJ membrane tortuosity and NM2B organization by recruiting cingulin to TJ. A branched network of actin filaments is associated with NM2B in the juxta-membrane region of apical junctions [12], and we propose that this branched network comprises γ-actin, and its tethering to TJ requires cingulin (Figure 6, Figure S6). Cytoplasmic γ-actin is not critical for junction assembly and organism viability [50, 51], but is specifically associated with the apical surface and TJ in epithelia [13, 14]. Labeling for γ-actin was reduced and shifted farther away from the membrane in cingulin-KO cells, and phalloidin labeling was decreased despite normal labeling of junctional β-actin. Since actin assembly at junctions is regulated by tension [52], our results suggest that when cingulin tethers to TJ the NM2B-γ-actin lace-like web, this latter is maintained under tension, and phalloidin binds very well to it. The KO of CGN uncouples the network from TJ and releases the tension, inducing the disassembly, and possibly the partial collapse of the network onto the peri-junctional ring, and decreased phalloidin labeling. Since the accumulation of NM2B is not abolished, but only altered/reduced in cells depleted of either ZO-1 [20, 21, 23] or cingulin, redundant mechanisms must exist to recruit NM2B to TJ and ZA. The mechanism through which cingulin tethers the γ-actin-NM2B network to TJ could be either through direct or indirect binding to myosin [32] or through recruitment of ARHGEF18 [39], or through modulation of ZO-1 conformation (see below), or a combination of these. Future studies should test these hypotheses and determine whether and how γ-actin is connected to the ZA. Although we did not observe any effect of cingulin KO on the localization of either NM2A or β-actin in our models, depletion of cingulin in human corneal epithelial (HCE) increases labeling for NM2A [39]. Possible reasons for this apparent discrepancy include the difference in cytoskeletal organization in stratified versus columnar/cuboidal epithelial cells, and the presence of residual cingulin in depleted cells, which could result in altered signaling through regulators of Rho family GTPases.

Actin and myosin control cellular mechanics, and here we studied how KO of CGN, CGNL1 or both affected apical surface stiffness and apicolateral membrane tension. In our atomic force microscopy experiments, the elastic modulus of WT MDCK cell is consistent with previously reported MDCK stiffness [53]. The loss of cingulin, and to a lesser extent of paracingulin, reduces the cell layer elastic stiffness, while monolayer integrity and junctions are maintained. This phenotype can be explained by a role of the branched γ-actin-NM2B network in providing mechanical support to the apical surface, consistent with γ-actin localization [13], and with the observed collapse of the apical mesh-like adhesion structure in cells lining the spinal canal, upon KO of NM2B in mice [54, 55]. Nonmuscle myosin isoforms have different enzymatic properties and cellular roles [56], and the longer actin-attachment lifetimes and greater strain dependence of NM2B allows it to maintain force more effectively than NM2A [57]. The loss of apical stiffness in CGN-KO cells correlates with a flatter shape of the apical membrane, in contrast to the phenotype of ZO-1-depleted cells, which show increased contractility and apical surface stiffness, and convex, extruding apical membranes [18, 19, 24]. Since ZO-1 recruits cingulin to TJ, we conclude that the complex cytoskeletal reorganization that occurs in ZO-1-depleted cells overrides the effects generated by the secondary loss of cingulin. This is not surprising, considering ZO-1’s multivalent interactions with proteins that bind to and regulate actomyosin filaments [58-60]. Using the FliptR membrane tension probe, we found that KO of cingulin, and to a lesser extent of paracingulin, increases overall apicolateral membrane tension, by reducing the sharp tension gradient existing in WT cells. This suggests that frictional forces between the membrane and the dense TJ-associated branched γ-actin network dampen the tension generated by the NM2A peri-junctional bundle, in agreement with previous observations [11, 12, 19, 61, 62]. Concerning the weaker mechanical phenotypes of paracingulin-KO cells, we speculate that similar mechanisms are involved, despite the apparently normal localizations of myosins, actins and ARHGEF18 in CGNL1-KO cells. The phenotypes may be weaker and harder to detect in CGNL1-KO cells because of the lower levels of paracingulin versus cingulin in epithelial cells [63], the small fraction of paracingulin associated with TJ versus ZA [30, 42] and the lower affinity of interaction of paracingulin with ZO-1. However, our experiments in different cell types, and using chimeric molecules to rescue NM2B localization indicate that cingulin and not paracingulin is specifically involved in the junctional assembly of NM2B. In contrast, both proteins were reported to regulate ARHGEF18, although in distinct cellular contexts [22, 39]. Thus, future studies should address the role of ARHGEF18 in membrane mechanics and actin and myosin isoform localization.

We show that the ZU5 domain, a 110-residue domain positioned at the extreme C-terminus of ZO-1 [5], binds to cingulin and paracingulin. Known interactors of the ZU5 domain of ZO-1 are the Cdc42 effector kinase MRCKβ, which is associated with actin at the leading edge of migrating cells, but is not localized at TJ [64, 65], and the Rho GTP Exchange factor ARHGEF11, which is recruited to TJ by ZO-1, but is not required for ZO-1 TJ targeting [58]. The ZU5-cingulin interaction not only recruits cingulin to TJ, but also promotes the unfolding of ZO-1, as operationally defined by the rescue of junctional DbpA labeling in ZO-2-depleted cells [25]. This suggests that loss of cingulin phenocopies the effect of blebbistatin, and ZO-1 folding could thus be due to reduced action of NM2B with ZO-1. However, our experiments in vitro using the CGN(1-70) fragment, which is unlikely to recruit actomyosin, show that its binding to the ZU5 is sufficient to induce ZO-1-DbpA interaction, suggesting that the CGN(1-70) fragment unfolds ZO-1 by antagonizing ZU5-dependent intramolecular interactions. Thus, although ZO-1 obviously unfolds and forms condensates in the absence of cingulin, cingulin binding might regulate a pool of folded ZO-1, lower its critical concentration for liquid phase separation, and/or increase the kinetic of formation and the size of ZO-1 condensates. Moreover, our analysis of ZO-1 condensates in either WT or CGN-KO backgrounds indicates that one function of the ZU5 domain could be to inhibit the binding of F-actin to the ABR domain, and binding of cingulin could relieve this inhibition (Figure S6). The idea that the ZU5 domain might interact with upstream, ABR-containing sequences of the C-terminal half of ZO-1 is consistent with the observation that the binding of CGN(1-70) to the isolated ZU5 domain is stronger than to constructs encompassing larger C-terminal fragments (Figure 3A, Figure S3D). There is also evidence from FRAP studies that the ABR domain is involved in intramolecular ZO-1 interactions [28]. Moreover, the ABR lies within a disordered region of the C-terminal half of ZO-1 and shows low affinity of binding to actin [66], suggesting that conformational changes driven by ligand-ZU5 interactions may promote the ABR interaction with actin filaments. These hypotheses need to be tested by future studies.

The role of cingulin in promoting the efficient accumulation of ZO-1 at TJ provides a mechanistic explanation for the observed ZU5-dependent stabilization of ZO-1 at junctions [67, 68]. As noted above, this stabilization could be due to cingulin promoting the unfolding and phase separation of an otherwise soluble pool of ZO-1, by relieving a ZU5-dependent intramolecular inhibition which modulates the binding of actin filaments to the ABR. Finally, tension is required for TJ formation [69, 70] and cingulin may promote tension-dependent accumulation of ZO-1 by recruiting to junctions ARHGEF18, which activates junctional RhoA [39]. Interestingly, KO of cingulin but not paracingulin reduced junctional ARHGEF18 in our epithelial cell models, although an interaction of ARHGEF18 with paracingulin was reported in endothelial cells [22]. Analysis of condensates also indicates that ZO-1 can associate with ARHGEF18 independently of cingulin. Since ZO-1 junctional levels are only reduced, but not abolished by KO of CGN, other mechanisms which unfold/stretch ZO-1 are sufficient to allow most of ZO-1 to assemble at junctions in the absence of cingulin, in agreement with previous data [71, 72]. These other mechanisms include heterodimerization, actomyosin tension applied indirectly, through ZO-1-interactors, and phosphorylation (Figure S6), whereas the role of cingulin is to unfold the ZU5 domain, promote the complete TJ assembly of ZO-1, and tether it to the NM2B-γ-actin network.

We also provide evidence indicating that in addition to ZO-1 [26], cingulin and paracingulin undergo phase separation, at least when expressed at high concentrations in cells. In a previous study, we showed that overexpressed GFP-tagged cingulin and paracingulin form structures that undergo dynamic fusions and relaxations, disappear upon mitosis, and rapidly associate with cell-cell contacts immediately following cytokinesis [49]. Cingulin and paracingulin contain intrinsically disordered N-terminal head and C-terminal tail domains, and a coiled-coil domain, all of which could drive phase separation, by enabling multivalent homo- and heterotypic protein-protein interactions [73, 74]. Furthermore, both cingulin and paracingulin are expected to form condensates based on prediction algorhythms [75]. Although additional work is required to characterize in detail the biophysics and biochemistry of cingulin and paracingulin condensates and establish their physiological relevance, here we show that ZO-1 condensates recruit cingulin and paracingulin, but cingulin and paracingulin condensates do not recruit ZO-1, in agreement with previous data [22, 39-41, 71, 72].

In summary, cingulin and paracingulin regulate the mechanical properties of the apical surface and the apicolateral plasma membrane, and cingulin regulates the conformation and TJ assembly of ZO-1 by organizing NM2B and γ-actin at the TJ-apical complex. Importantly, both cingulin and paracingulin associate with microtubules, and cingulin organizes the epithelial planar apical network of microtubules [6, 35, 76, 77]. Thus, the present results suggest that cingulin and paracingulin are major players in the cross-talk between the microtubule and actin cytoskeletons at junctions and apical membranes, to fine-tune the mechano-regulation of morphogenesis, TJ barrier function, and other physiological and pathological aspects of epithelial and endothelial cells.

## ACKNOWLEDGEMENTS

This study was supported by the Swiss National Fund for Scientific Research (N. 31003A_152899 and N. 31003A_172809 to S.C., N. 31003A_130520, N.31003A_149975 and N.31003A_173087 to A.R.), European Research Council Consolidator Grant N° 311536 (to AR). W.H. and J. Y. are funded by the Singapore Ministry of Education Academic Research through the MOE Research Scholarship Block (RSB) scheme (to W. H.) and the Singapore Ministry of Education under the Research Centres of Excellence program (to J. Y.). We thank Jerôme Bosset and Christoph Bauer (BioImaging Facility, Faculty of Sciences) for help with scanning electron microscopy, the cited colleagues for gifts of reagents, and David Shore and Jean-Claude Martinou for comments on the manuscript.

## AUTHOR CONTRIBUTIONS

Conceptualization, S.C., E.V., F.R., D.S.; Methodology, E.V., F.R., D.S., W.H., A.C., A.F., J.S., and S.C.; Validation, E.V., F.R., D.S., W.H., A.C., A.F., J.S., and S.C.; Formal Analysis, E.V., F.R., D.S., W.H., and A.C.; Investigation, E.V., F.R., D.S., W.H., A.C., A.F., J.S., I.M., L.J., and S.C.; Resources, E.V., F.R., D.S., V.D., C.C., S.S., I.M., L.J., and S.C.; Data Curation, E.V., F.R., D.S., W.H., A.C., A.F., J.S. I.M., and L.J.; Writing – Original Draft, S.C., E.V., F.R., D.S., W.H., A.C., A.R., J.Y; Writing – Review & Editing, S.C., E.V., F.R., W.H., A.C., A.F., J.S., V.D., C.C., S.S., A.R., J.Y.; Visualization, S.C., E.V., F.R., D.S., W.H., A.C., A.F.; Supervision, S.C., J. Y. and A.R..; Project Administration, S.C.; Funding Acquisition, S.C., J. Y. and A.R..

## STAR ★Methods

### Resource Availability

#### Lead Contact

Further information and requests for resources and reagents should be directed to and will be fulfilled by the Lead Contact, Sandra Citi.

#### Materials Availability

Reagents generated in this study will be made available on request, but we may require a payment for shipping and a completed Materials Transfer Agreement.

#### Data and Code Availability

This study did not generate any unique datasets or code.

### Experimental Model and Subject Details

Eph4 (mouse mammary epithelial cell line) WT and ZO-1-KO cells, MDCK (Madin-Darby Canine Kidney – II), mCCD (mouse cortical collecting duct epithelial cell line) and HEK293T cells were cultured at 37°C, 5% CO_2_ in DMEM containing 10% or 20% FBS (for mCCD). For Eph4, MDCK and mCCD culture media were supplemented with 1% non-essential amino acids, 100 units/ml penicillin and 100 μg/ml streptomycin [25, 63, 78]. The sex of Eph4 and MDCK lines was female. For other lines, sex was not listed in the information available to us. We trusted the providers of the cells for their authentication.

Cell lines KO for cingulin (CGN) and paracingulin (CGNL1) were generated using CRISPR/Cas9 gene editing technology, designing guide-RNA (gRNA) using the Zhang Lab CRISPR design tool, targeting exons which are present in all major transcripts of CGN and CGNL1. For mouse CGN and CGNL1 (mCGN, mCGNL1) the target sequences for the CRISPR/Cas9, to generate Eph4 and mCCD KO lines, were selected in exon 2 (Key Resource Table). For canine CGN and CGNL1 (cCGN, cCGNL1) the target sequences for the CRISPR/Cas9 for MDCK cells were selected in exon 1 (Key Resource Table). The gRNAs were subcloned into the BbsI site of pSpCas9(BB)-2A-GFP (PX458) CRISPR plasmid. Cells were transfected using Lipofectamine 2000. At 48h post-transfection, single cells were sorted (using a Beckman Coulter MoFlo Astrios sorter, Flow Cytometry Service, University of Geneva Medical School) into 96-well tissue culture plates. Single clones were further amplified and screened for the KO using immunoblot and immunofluorescence analyses, based on which two-three CGN-KO and CGNL1-KO clones (one CGNL1-KO clone for Eph4) were selected and verified by sequencing. For sequencing, genomic DNA was extracted using DNeasy Blood & Tissue kit, the genomic loci of CGN and CGNL1 (around 500 bp upstream and 600 downstream of target sequence) were amplified by PCR using specific primers (Key Resource Table) and subcloned into pBluescript II KS(+) using restriction enzymes SacII and XhoI for mCGN, XbaI and EcoRI for mCGNL1 and cCGN, BamHI and SalI for cCGNL1. The T7 primer was used for sequencing inserts for genotyping.

To generate CGN/CGNL1 double-KO MDCK clones, CGN-KO (to generate clone 21D3) or CGNL1-KO (to generate clone 11C9) clones were transfected with above-mentioned CGNL1-KO or CGN-KO CRISPR/Cas9 constructs, respectively, sorted, amplified and screened as described above for single KO.

To generate MDCK CGNL1-KO cells stably expressing YFP-myc, cells were transfected using Lipofectamine 2000. At 48h post-transfection, single cells were sorted into 96-well tissue culture plates and were grown in the presence of 200 µg/ml Hygromycin B prior to clone amplification.

### Method Details

#### Antibodies and immunofluorescence

Antibodies are described in Key Resources Table.

For immunofluorescence (IF) cells were cultured either on glass coverslips in 24-well plates for 3 days seeded at a density of 1-2 x 10^5^ cells/well (Figure 3; Figure 4; Figure S1; Figure S2 (B, C, D, G, H, I, J); Figure S3 (H, I); Figure S4) or on 24-mm Transwell filters for 5 days seeded at a density of 5 x 10^5^ cells (Figure 2; Figure S2 (E, F, K, L)).

IF for cells grown on coverslips was carried out by washing cells 2x with cold PBS, fixing in methanol at -20° for 10 min, washing 3×5 min with PBS, incubating with primary antibody (either at RT for 1 h, or for 16h at 4°C), followed by washing 3x with PBS, incubating with secondary antibody (30 min at 37°C in a humidified chamber), washing 3x with PBS, and mounting either with Vectashield with DAPI or Fluoromount-G.

For IF of actins (F-actin-phalloidin, β-actin, γ-actin) in Eph4 cells, cells were fixed in room temperature 1% PFA for 5 min, followed by rinsing 2x with PBS and incubating with methanol at -20°C for 5 min, followed by gradual rehydration in PBS (3x replacing 50% of volume with PBS), 2x washes in PBS [13, 14], and mounting with Fluoromount-G.

For IF of DbpA, Eph4 cells were permeabilized, fixed and labelled as described in [25].

IF analysis of CGN, β-catenin and occludin in MDCK cells grown on Transwell, cells were washed 2x with cold PBS, fixed in methanol for 16h at -20°C), followed by 1-min treatment with acetone (−20°C). Filters were excised manually using a razor blade and hydrated in IMF buffer (0.1% Triton X-100, 0.15 M NaCl, 5 mM EDTA, 20 mM HEPES, pH 7.5, 0.02% NaN_3_ as preservative).

For IF analysis of NM2A, NM2B, F-actin-phalloidin and ARHGEF18 in MDCK cells grown on Transwells, cells were washed with PBS containing Ca^2+^ and Mg^2+^ (PBS++), fixed in 1% PFA pre-warmed at 30°C for 7 min, followed by rinsing 2x with PBS++ and incubating with methanol at -20°C for 5 min, followed by gradual rehydration in PBS++ (3x replacing 50% of volume with PBS), and 2x washes in PBS++ [13, 14]. Cells were permeabilized with 0.2% of Tx100 in PBS++ 15min at RT and saturated 30 min with 2% of BSA in PBS++.

For mCCD cells grown on Transwells, fixation for IF of NM2A, NM2B and actins was carried out using the same protocol above, except that final permeabilization and blocking steps were carried out together, using PBS++ containing 0.2% Tx100 and 2% BSA (30 min at RT).

Incubation with primary antibodies was carried out for 16h at 4°C (in a humidified chamber) and with secondary antibodies for 1-2h at RT. The filters were placed on glass slides with the cells facing up and were mounted with Fluoromount-G.

Slides were imaged on a Zeiss LSM800 confocal microscope using a Plan-Apochromat 63x/1.40 oil objective at a resolution of 1024×1024 px or on an upright microscope Leica DM4B (Figure S1 (E-G, K-L); Figure S2 (I)) using 63x oil objective at a resolution of 2048×2048 px; the pixel size=0.10 μm. Single confocal plane images are shown in Figure 2D; Figure 3I; Figure 4; Figure S1 (M, Q); Figure S2 (B-D), Figure S4. Maximum intensity projections of z-stack images are shown in Figure 2A (6 confocal planes over 8 µm, step size=1.6 µm); Figure 2B (5 confocal planes over 1 µm, step size=0.25 µm); Figure 2 (F-L) and Figure S2 (E-F) (4 confocal planes over 1.2 µm, step size=0.4 µm); Figures 2(M-N) and Figure S2 (K-L) (5-6 confocal planes over 1-1.3 µm, step size=0.25 µm); Figure 3D (7 confocal planes over 6 µm, step size=1 µm; Figure 3 (G-H) (4 confocal planes over 3 µm, step size=1 µm); Figure S2 (G-H, J) (5 confocal planes over 1.6 µm, step size=0.35 µm) Figure S3 (H, I) (4-5 confocal planes over 1.1-1.4 µm, step size=0.4 µm. Images were extracted from .lif, .lsm or .czi files using ImageJ, adjusted and cropped using Adobe Photoshop, and assembled in Adobe Illustrator figures.

#### Measurement of cell proliferation

To measure cell proliferation, cells were plated in 12-well plate (75 000 cells/well) in duplicate, and cells in one well were trypsinized and counted in a hemocytometer each day, for 6 days.

#### Plasmids

Full length (FL) mCGN (S2407) and mutant lacking ZIM domain (Δ1-70) (S2408) were generated by PCR using appropriate oligonucleotides containing Kozak sequence and subcloning into XhoI-KpnI sites of pcDNA3.1(-)/MycHis C. GFP-myc (S1166) construct was generated by subcloning of GFP-myc into pcDNA3.1(-)/MycHis A. The FL mCGN (S2363) and mCGNL1 (S2386) constructs were generated by PCR and subcloning into NotI-ClaI sites of pTRE2Hyg containing GFP and myc tags, downstream of GFP and upstream of myc. GFP-tagged human (h) CGN FL (S2508) (G-CGN-FL) and the mutant lacking ZIM domain (G-CGN-Δ1-70) (S2509) were generated by PCR and subcloning into NotI-Acc65I sites of GFP-myc (S1166) downstream of GFP.

FL cCGN (S1136) and FL cCGNL1 (S2432) HA-tagged in C-terminal were generated by PCR using appropriate oligonucleotides containing Kozak sequence and subcloning into BamHI-KpnI sites of pcDNA3.1(-)/MycHis A and BamHI-NotI sites of pcDNA3.1/Zeo(+), respectively. Chimeras of cCGN and cCGNL1 HA-tagged in C-terminal (S2433: aa 1-579 from CGNL1 and 339-1190 from CGN; S2434: aa 1-338 from CGN and 580-1295 from CGNL1) were obtained by triple PCR using appropriate oligonucleotides containing Kozak sequence and overlapping region for fusion between the head and rod-tail sequences, before insertion into BamHI-NotI sites of pcDNA3.1/Zeo(+).

FL cCGN (S2294=S1052) was described previously [79]. YFP-tagged cCGNL1 (S1137) and YFP-myc (S1152) were described previously [34]. GST (S1851) and GFP (S1821) tagged CGN (1-70aa) were constructed by PCR on cCGN and subcloned either into pGEX4T1 (EcoRI-NotI) or into GFP-containing pTRE2-hyg (S1210) [80] downstream of GFP (BglII-NotI), respectively. The FL myc-hZO-1-HA (S1947) construct; GFP-tagged constructs of fragments within the C-terminal region of hZO-1 (residues 888-1619 (S1807); 1150-1619 (S1808); 888-1748 (S1809); 1150-1748 (S1810); 1619-1748 (S1811)); and FL hZO-1 for expression in insect cells were described previously [25]. GFP-tagged constructs of fragments within the C-terminal region of hZO-1 (residues 1698-1748 (S1874)) and (residues 1550-1650 (S1875)) were generated by PCR and subcloned downstream of GFP into NotI-KpnI sites of pCDNA3.1(+). The construct of myc-hZO-1-HA (S2161) lacking the ZU5 domain (ZO1-ΔZU5, residues 1-1619) and GFP tagged mZO-1 (S2474) were described previously [78]. GST-DbpA, CFP-HA and HA-DbpA were previously described [47]. The DbpA-binding construct of GST fused to the C1 domain of GEF-H1 was a kind gift from K. Matter [48]. FL hCGN (S2411), FL hCGNL1 (S2442) (CGNL1-FL-HA) and mutant lacking ZIM domain (S2510) (CGNL1-Δ1-80-HA) were generated by PCR using appropriate oligonucleotides containing Kozak sequence and subcloning upstream of HA into modified HA containing pCDNA3.1 (BamHI-XbaI sites for S2411, BamHI-NotI sites for S2442 and S2510).

GST-tagged hZU5 (S1789), ZIM domain – containing fragments of hCGN (1-121aa) (S96) and (1-353aa) (S97) and hCGNL1 (1-119aa) (S1024) were generated by PCR and subcloning into pGEX4T1 (EcoRI-XhoI sites for S1789, S96, S97; BamHI-EcoRI sites for S1024).

#### Recombinant Protein Expression, Glutathione S-Transferase (GST) Pulldown, and Supernatant Depletion Assays

GST fusion proteins were expressed in BL21 bacteria and purified by affinity chromatography on Glutathione Sepharose [47] or magnetic beads. Pulldowns were carried out using lysates either from insect cells expressing FL ZO-1, or from lysates of HEK293T cells expressing either GFP-tagged ZO-1 fragments; GFP-tagged CGN constructs, HA-tagged DbpA and CGNL1 constructs (prey) as previously described [78, 81]. Lysates containing prey proteins were normalized, and equivalent amounts of prey proteins were added to 5 µg of baits (GST tagged DbpA, CGN fragments (residues: 1-70, 1-121, 1-353), CGNL1 (1-119), ZU-5 domain of ZO-1, C1 domain of GEF-H1).

To determine the K_d_ of interaction between the ZU5 domain of ZO-1 (1619-1748=Cter) and the CGN1-70 fusion protein, we used a quantitative GST-pulldown Supernatant Depletion Assay [25, 47]. Increasing volumes (1, 2, 5, 10, 12.5, 15, 17.5, and 20 μl) of Glutathione Sepharose beads pre-loaded with GST-CGN(1-70) bait (0.666 μM), were added to prey protein (GFP-ZO-1-C-ter), in a total volume of 0.5 ml CO-IP buffer. As a negative control, beads pre-incubated with CO-IP buffer (1, 2, 5, 10 μl) were added to the prey. After 16h incubation at 4°C, beads were pelleted at 16,000 x *g*, and the prey proteins remaining in the supernatant were analyzed by SDS-PAGE and immunoblotting. GST pulldown experiments to assess the effect of CGN(1-70) on the interaction between DbpA and full-length ZO-1 were carried out by incubating GST-DbpA (5 μg) with insect cell lysate containing full-length ZO-1, in the presence of increasing volumes of normalized HEK293T cells lysates containing either GFP-CGN(1-70) or GFP, this latter as a negative control. Although it is unlikely that sufficient concentrations of additional interacting partners of baits and preys are present in the GST pulldown assays and might influence some of the results, this possibility cannot be formally excluded. Concentration of recombinant proteins was determined by densitometric analysis of Coomassie-stained SDS gels, compared to a BSA standard curve.

#### Transfection, siRNA-mediated ZO-2 depletion and exogenous expression of proteins

For transfections for rescue experiments, cells were plated on glass round 12 mm coverslips in 24-well plate (100 000 cells/well), transfected next day using Lipofectamine 2000 or jetOPTIMUS DNA transfection reagent and fixed for IF 72 h post-transfection.

For transfections of cells grown on Transwells cells were transfected 24h after plating using jetOPTIMUS according to the manufacturer’s protocol and fixed for IF 4 days after transfection.

For siRNA-mediated ZO-2 depletion (target sequence in Key Resource Table) cells were transfected with Lipofectamine RNAiMAX 24h after plating, using siRNA negative control as negative control.

For siRNA and DNA co-transfections, Eph4 cells were transfected next day after plating with the mix of siRNA and DNA using Lipofectamine 2000, 48h post-transfection cells were treated with 50 μM blebbistatin for 16 h and then fixed [25].

HEK cells were plated in 10 cm dish (2×10^6^ cells/dish), transfected next day using Lipofectamine 2000 and lysed 48h post-transfection.

#### Overexpression and condensate analysis, live microscopy

To study biomolecular condensates of CGN, ZO-1 and CGNL1 (Figure 5 (A, B) and Figure S5A respectively), 5×10^5^ MDCKII cells were seeded onto 35-mm glass bottom culture dishes. 24h after plating cells were transfected with 8 µg of DNA per dish using JetOptimus®. GFP-CGN or GFP-CGNL1 or GFP-ZO1 condensates were analyzed 48h post transfection. Live imaging was performed on a Zeiss LSM780 confocal microscope using argon laser with the 488 nm excitation and PlanApo N60x oil /1.4 objective. Single plane images were recorded every 2s for 100s at a resolution of 512×512 px.

To study recruitment of client proteins by condensates, 1×10^5^ MDCKII cells were seeded onto glass coverslips placed in 24 well plates. 24h after plating cells were transfected with 1,5 µg of DNA per dish using JetOptimus®. 48h post transfection cells were fixed using protocol for NM2A for MDCKII on Transwells, described in immunofluorescence section. Images were taken using Leica DM4B (Figures 3 (E-F); 5 (C-E); S5 (B-D)) or Zeiss LSM800 for phalloidin (Figures 5 (C-G); S5 (C-D)). Maximum intensity projections of four confocal planes representing 1.2 µm final depth are shown for phalloidin. Imaging settings and treatment were as described in immunofluorescence section.

#### Scanning Electron Microscopy

Confluent monolayers were rinsed with PBS and fixed with 2% glutaraldehyde for 20 min at RT, after washing with 0,1M sodium cacodylate buffer cells were post-fixed with 1% osmium tetraoxide in 0,1M sodium cacodylate buffer for 20 min at RT. Cells were rinsed with sodium cacodylate buffer and water and then dehydrated with a series of ethanol washes (30, 50, 70, 95 and 100%), followed by acetone washes. Samples were dried using critical point dryer (Leica EM CPD030), coated with gold and observed under the JEOL-6510LV low vacuum scanning electron microscope.

#### Immunoblotting

Cell lysates were obtained using RIPA buffer containing 1X cOmplete or Pierce protease inhibitor and immonoblotting was performed as previously described [25, Vasileva, 2017 #4538]. β-tubulin was used for protein loading normalization in immunoblots. Numbers on the left of immunoblots indicate migration of molecular size markers (kDa).

#### Atomic force microscopy indentation measurements

The AFM-based indentation measurements were carried out using a commercial AFM (Dimension FastScan, Icon Scanner, Bruker). A polystyrene bead (5 µm radius, Invitrogen) was stuck on the tipless silicon nitride cantilever (MLCT-O10-E, Bruker) by epoxy fix. The spring constant of the home-made cantilevers, calibrated each time before measurement by thermal fluctuation method, were in the range of 0.10∼0.15 N m^-1^.

All AFM indentation measurements were realized in cell culture medium at room temperature. The cells were cultured in 60 mm petri dish for 36 h in incubator until forming monolayers (confluency > 80%). In a typical experiment, the cantilever was brought to the cell layer with the constant speed of 1 µm s^-1^ until reaching the maximum contact force of 5 nN, where the maximum indentation distance of cells was in the range of 0.5-1.5 µm. Then the cantilever was retracted and moved to another spot for the next cycle. A box pattern containing 100 spots in 40 µm × 40 µm region was set and typically 5-10 such regions were randomly selected in each measurement to obtain the averaged stiffness of the cell.

The force-indentation traces were analyzed to obtain the Young’s modulus of the cells by using the NanoScope Analysis program. After baseline correction and contact point estimation, the approaching force-indentation curve was fitted with the Hertz (Spherical) model (eq. 1) in the contact force range from 0.5 nN to 4.5 nN. Constant parameters and data range were chosen to minimize the bias for different cell types.

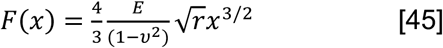

where F is the force of the cantilever, x is the indentation distance of the cell pressed by the cantilever, E is the Young’s modulus of the cell layer, r is the radius of the spherical indenter, and U is the Poisson ratio. The Poisson ratio of cell is normally in the range of 0.3-0.5. We chose U=0.5 in all calculations.

#### Lifetime measurements

Lifetime imaging was carried out on MDCK cells seeded on 35-mm glass bottom dishes at a density of 3×10^5^ cells/dish and grown for 4 days. 1,5 μl of FliptR solution was added to each dish prior imaging. FLIM imaging was performed as described [46].

FLIM imaging was performed using a Nikon Eclipse Ti A1R microscope equipped with a Time Correlated Single-Photon Counting module from PicoQuant [82]. Excitation was performed using a pulsed 485nm laser (PicoQuant, LDH-D-C-485) operating at 20 MHz, and emission signal was collected through a bandpass 600/50nm filter using a gated PMA hybrid 40 detector and a TimeHarp 260 PICO board (PicoQuant).

#### Quantification and Statistical Analysis

Data processing and analysis were performed in GraphPad Prism 8. All experiments were carried out at least in duplicate, and data are shown either as dot-plots, as histograms or as line-graph (with mean and standard deviation indicated). Statistical significance of quantitative data was determined by unpaired two-tailed Student’s t-test (when comparing two sets of data), ordinary one-or two-way ANOVA with Tukey’s post hoc test (for multiple comparisons), (ns=not significant difference)=p>0.05, *p≤0.5, **p≤0.01, ***p≤0.001,**** p≤0.0001).

#### Analysis of immunofluorescence data

For the quantification of junctional immunofluorescent signal (Figure 4; Figure S3; Figure S4), pixel intensity for each channel was measured in the selected junctional area using the polyhedral tool of ImageJ, and the averaged background signal of the image was subtracted. Relative intensity signal was expressed as a ratio between the signal of protein of interest and an internal junctional reference (either PLEKHA7 or occludin, or PLEKHA6). 9-45 junctional segments were analyzed.

For the measurement of the zig-zag index (L(TJ)/L(St)) (Figure 2 (C, E); Figure S2A): ratio between actual length of bicellular junction and the distance between two vertexes), we used the method described in [23], and measured the length of the TJ (L(TJ)) by using the freehand line trace in ImageJ, and the straight length of junction (L(St)) by using a straight line between vertexes. 45-240 junctions were analyzed.

To analyze intensity and distribution of β- and γ-actins the line across junction was drawn using Straight line tool in ImageJ and line scan graphs were generated by plotting profile. Full width at half-maximum (FWHM) pixel intensity was determined for actins and PLEKHA7, and the actin/PLEKHA7 ratio was calculated. 30 junctions were analyzed.

Each dot of dot-plot graphs represents one measurement, and data are shown in arbitrary units (a.u.).

#### Analysis of condensates

To analyze recruitment of F-actin (TRITC-Phalloidin) into ZO-1 condensates (Figure 5) ZO-1 condensates were encircled using Polygon selection tool in ImageJ, relative fluorescent intensities of ZO-1 and F-actin were measured, and F-actin/ZO-1 ratio was calculated after background signal subtraction.

Each dot of dot-plot graphs represents one measurement, and relative intensities are shown in arbitrary units (a.u.). 30 droplets were analyzed.

#### Analysis of immunoblotting data

For the quantification of immunoblot signals (Figure S3), densitometric analysis was carried out on band signals from at least 3 separate experiments. Each time the signal was normalized to tubulin, used as reference for total protein loading, and expressed as percentage taking the strongest signal as 100%.

#### Analysis of atomic force microscopy indentation measurements

Each dot of dot-plot graphs represents one Young’s modulus value from one indentation curve. Note that the indentation is site-by-site, not cell-by-cell, representing the local stiffness of the cells.

#### Analysis of lifetime measurements

SymPhoTime 64 software (PicoQuant) was used to fit fluorescence decay data (from full images or regions of interest) to a dual exponential model after deconvolution for the instrument response function (measured using the backscattered emission light of a 1 µM fluorescein solution with 4M KI). Data was expressed as means ± standard deviation of the mean. Slopes were determined based on simple linear regression model.

## KEY RESOURCES TABLE

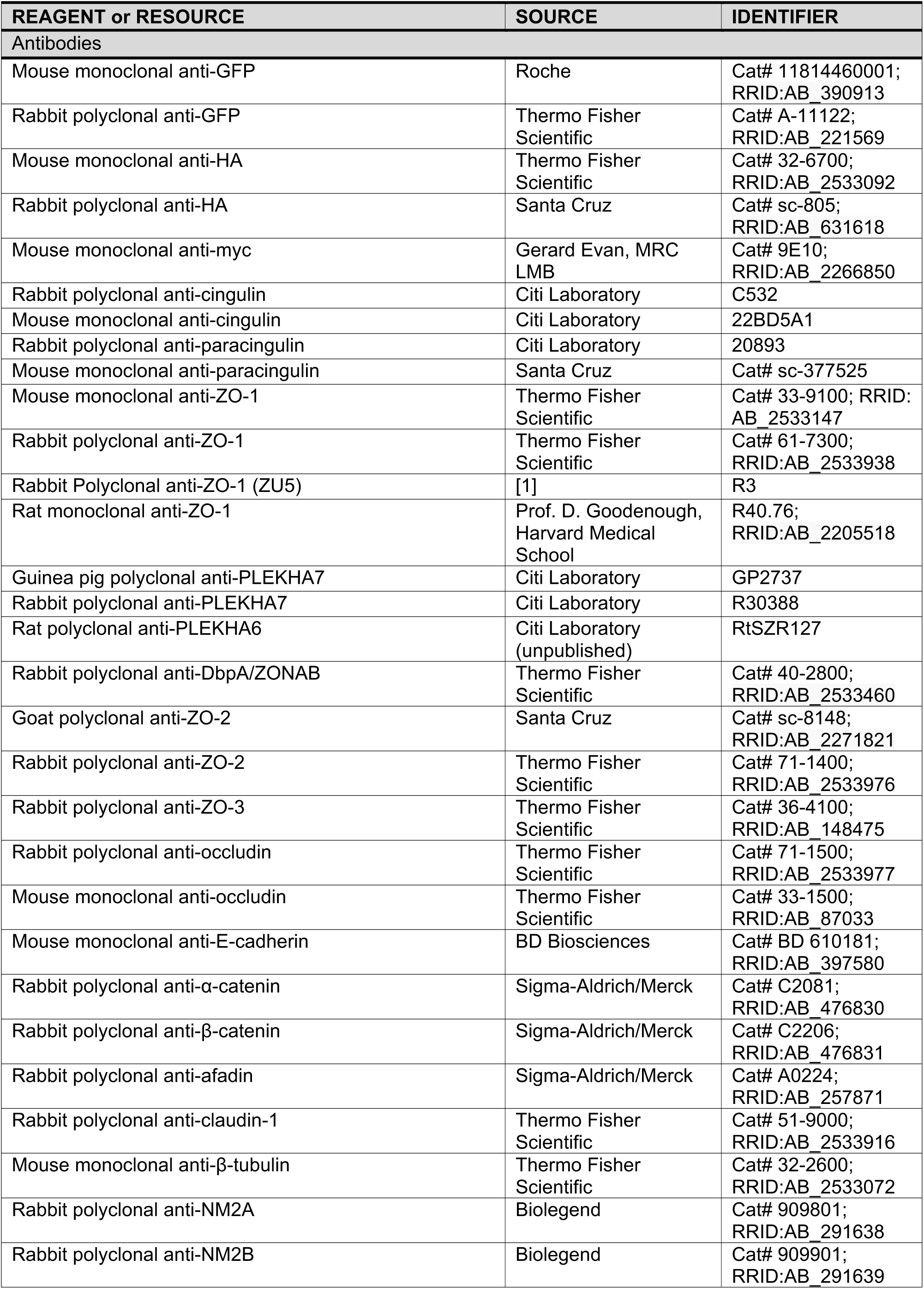
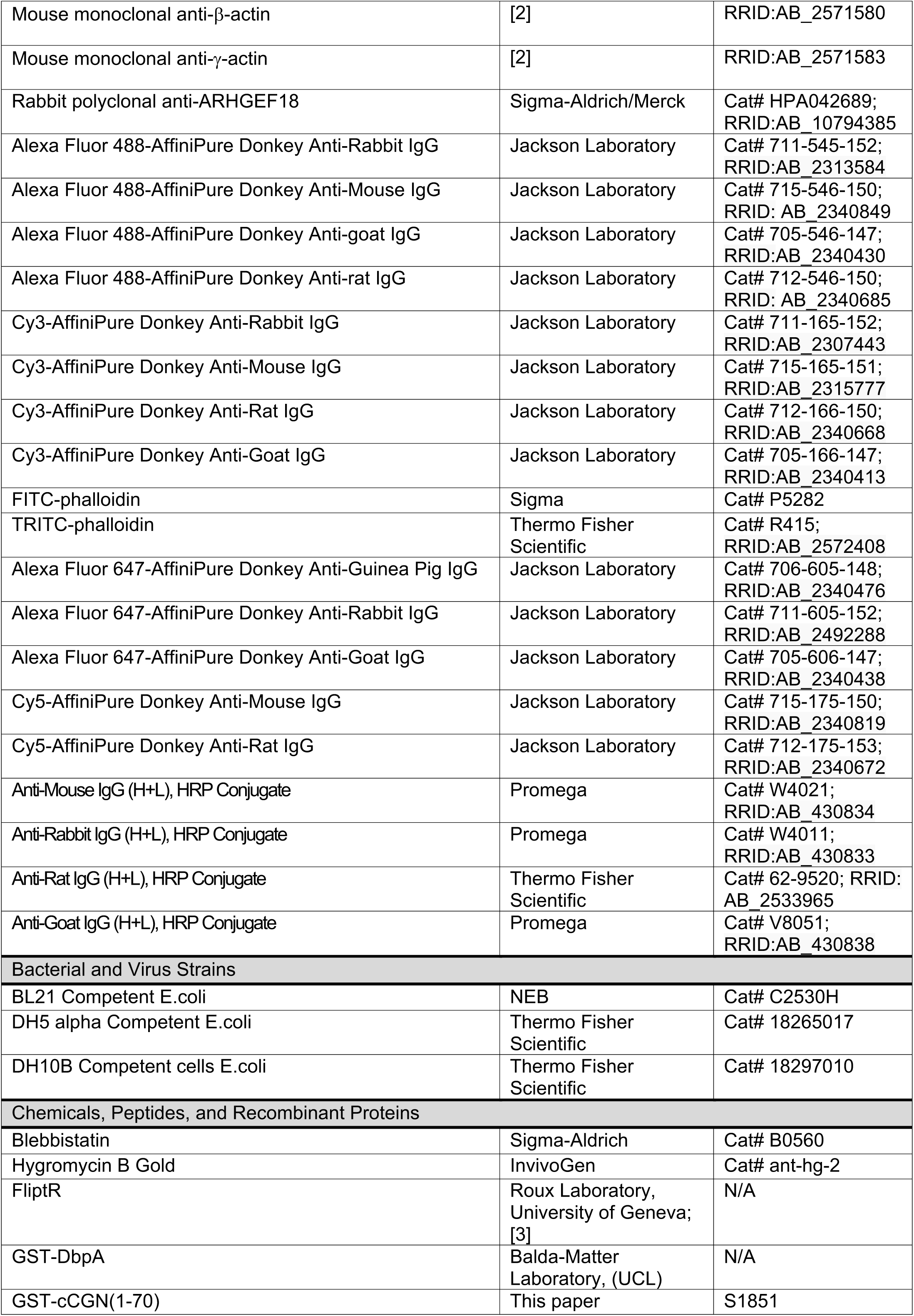
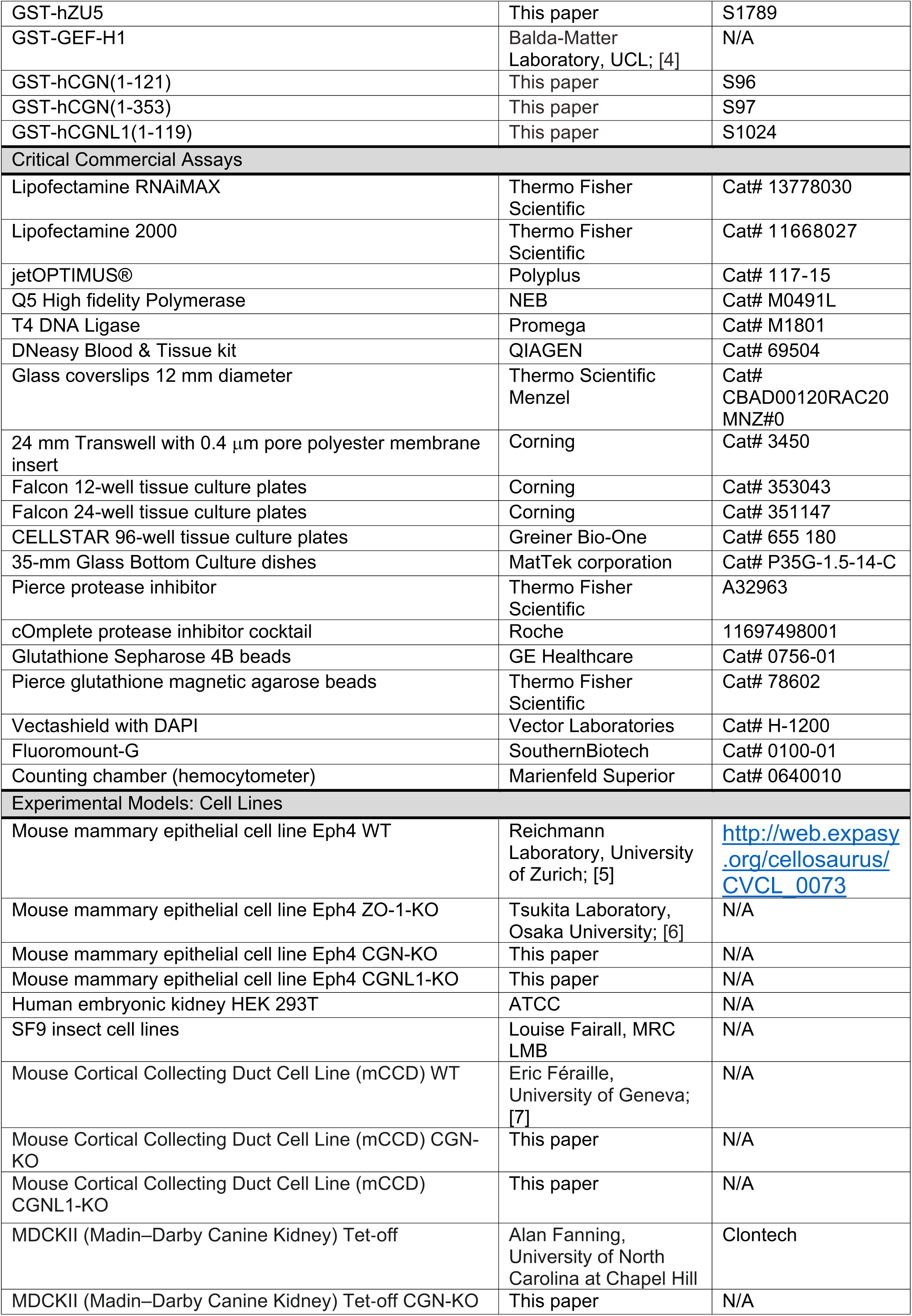
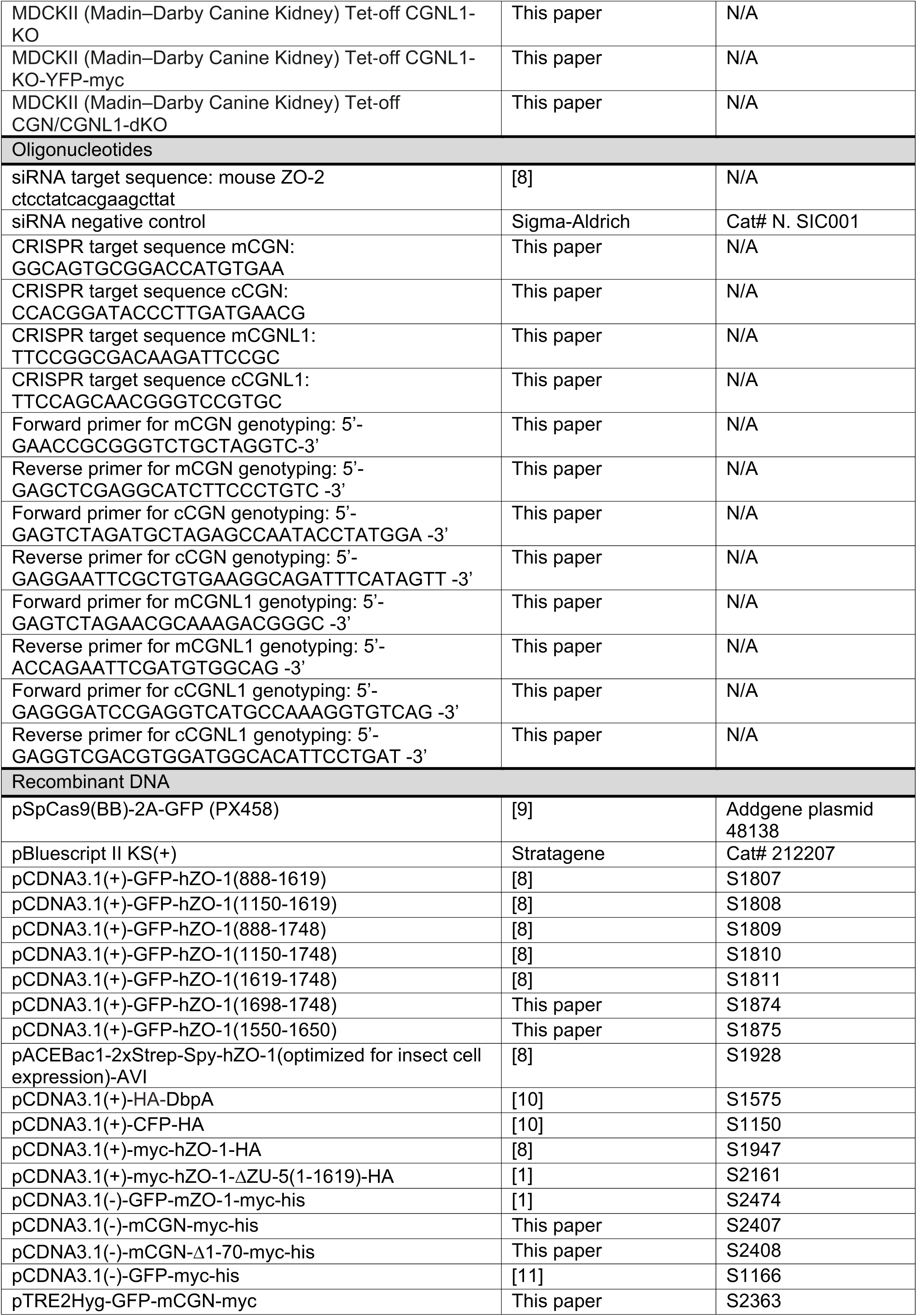
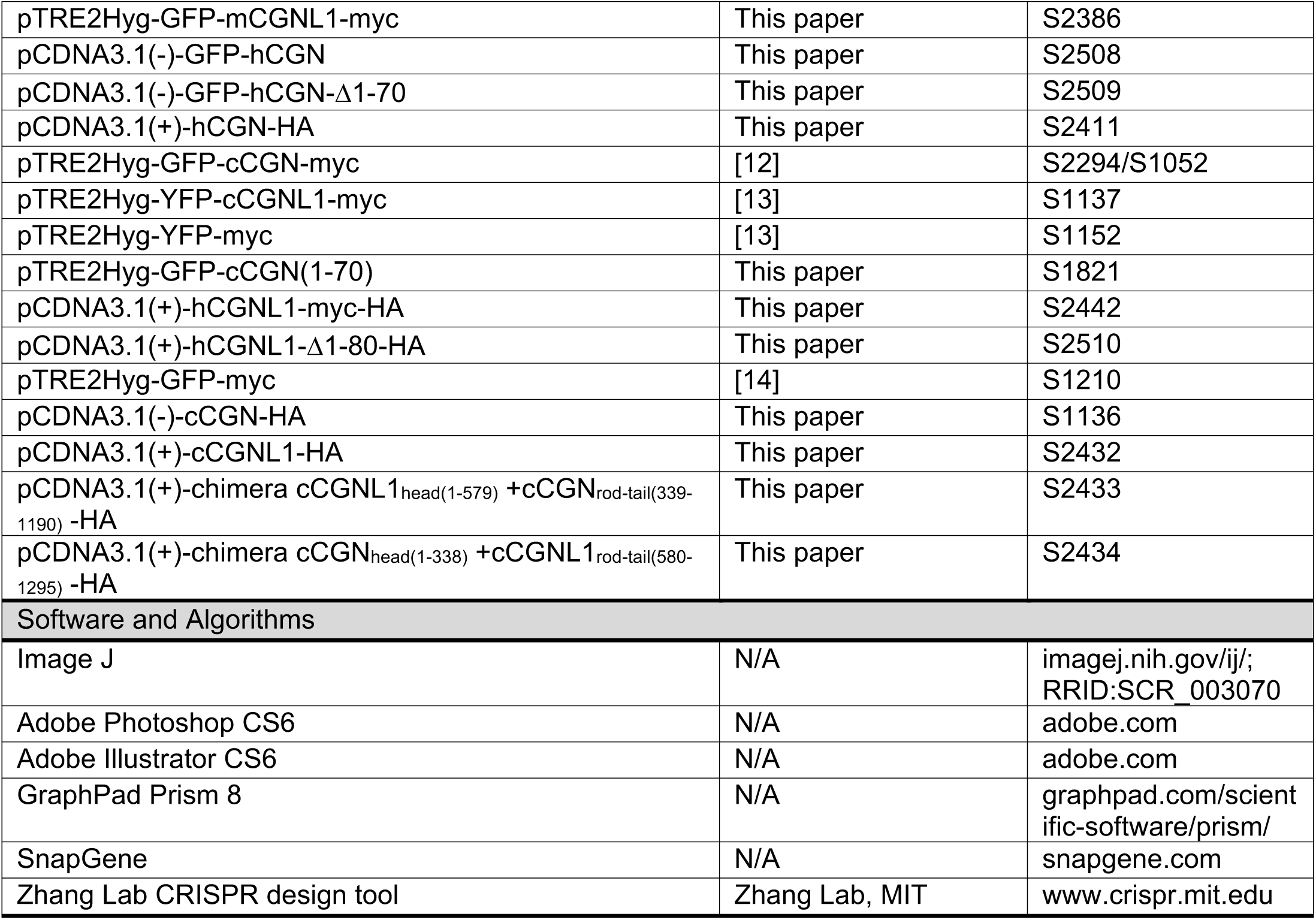

## Table

**Table S1.** Young’s modulus values for MDCK lines.

| Line genotype | Young’s modulus (MPa) |
| --- | --- |
| WT | 0.0037 ± 0.0014 |
| CGN-KO | 0.0015 ± 0.0007 |
| CGNL1-KO | 0.0021 ± 0.0019 |
| CGN/CGNL1-dKO | 0.0014 ± 0.0005 |

## Supplemental Data

**Figure S1.**
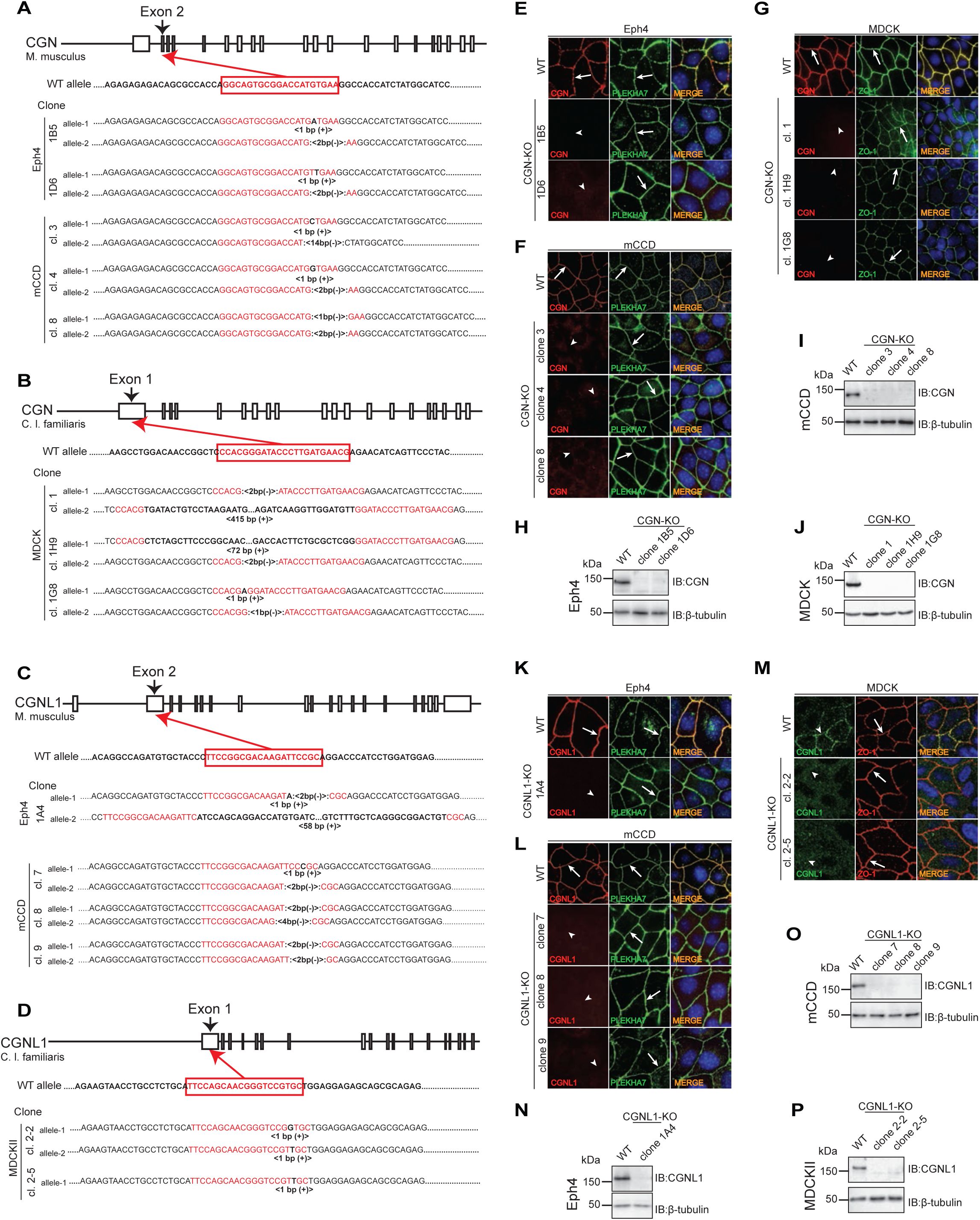
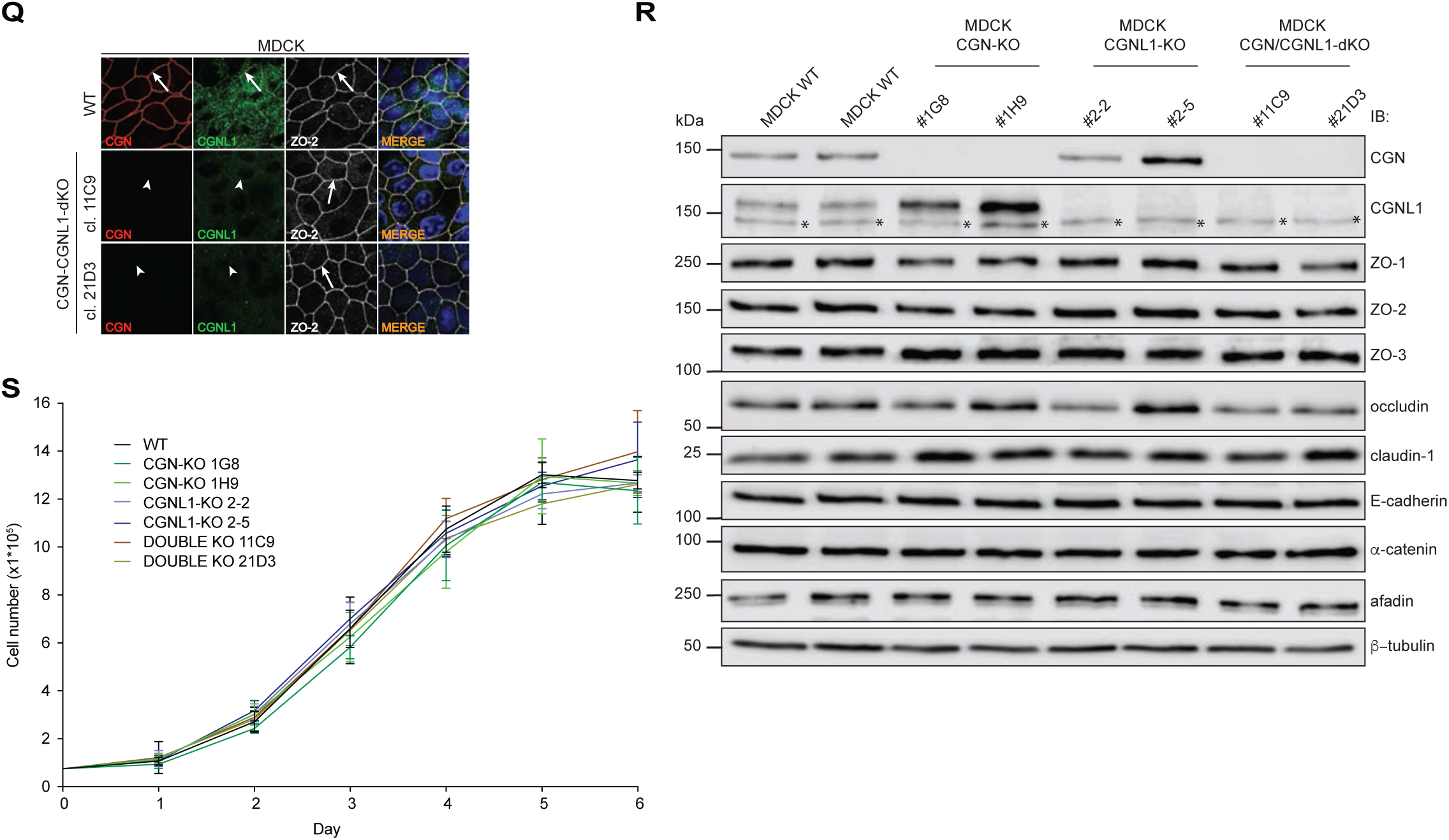
(Related to Figures 1-5). Generation and phenotypic characterization of knock-out (KO) clonal lines for either cingulin, paracingulin or both in Eph4, mCCD and MDCK cells. (A-D) Generation of CRISPR-KO clones of either CGN (A-B) or CGNL1 (C-D) in Eph4 (A, C), mCCD (A, C) and MDCK (B, D) cells. Each scheme shows exon-intron structure, WT alleles sequence, guide RNA target sequence (red), mutations, insertions (“+”) and deletions (“-”)(bold) in each allele. Number of inserted or deleted nucleotides are indicated for each allele. Sequencing of the second allele for MDCK CGNL1-KO clone 2-5 was not successful probably because of huge insertion since there were two bands on the gel after amplification of the target region. (E-J) Phenotypic characterization of CGN-KO clonal lines of Eph4 (E, H), mCCD (F, I) or MDCK (G, J) cells, either by immunofluorescence (E-G) or immunoblotting (I-J). Cells were labeled with antibodies against CGN and either PLEKHA7 or ZO-1, as reference junctional markers. Identifier for each clonal line is indicated on the left. Nuclei are stained with DAPI. Arrows indicate junctional localization, arrowheads indicate undetectable/decreased labeling. For immunoblots total RIPA lysates were used (STAR methods), and β-tubulin was used as a loading control. Numbers on the left indicate apparent size in kDa, based on the migration of prestained molecular weight markers. (K-P) Phenotypic characterization of CGNL1-KO clonal lines of Eph4 (K, N), mCCD (L, O) or MDCK (M,P) cells, either by immunofluorescence (K-M) or immunoblotting (N-P). (Q-R) Phenotypic characterization of double-KO (CGN+CGNL1) clonal lines of MDCK cells, either by immunofluorescence (Q) or immunoblotting (R). The asterisk in (R) indicates a non-specific polypeptide labelled by anti-CGNL1 antibodies. (S) Proliferation curves of clonal KO lines of MDCK cells.

**Figure S2.**
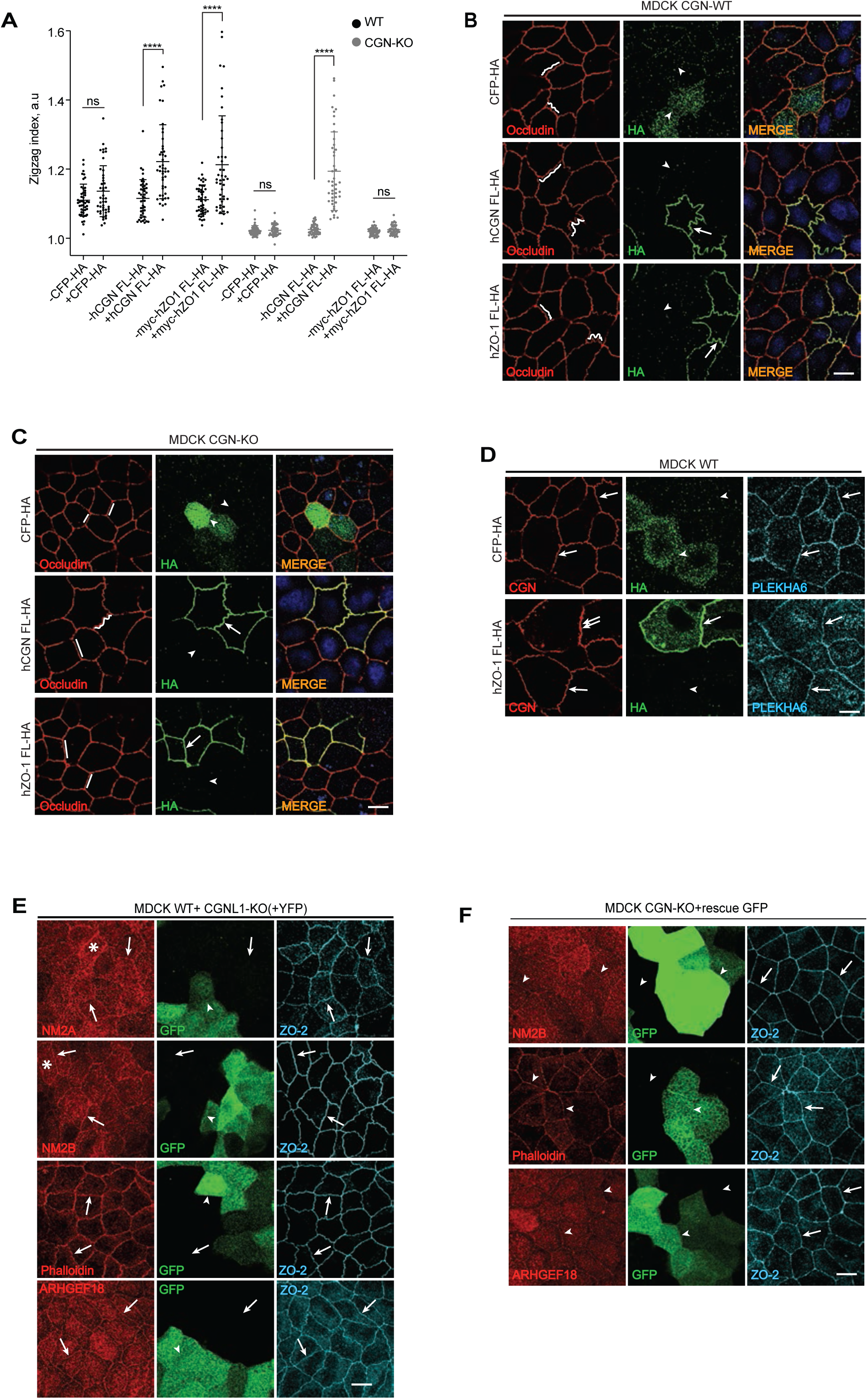
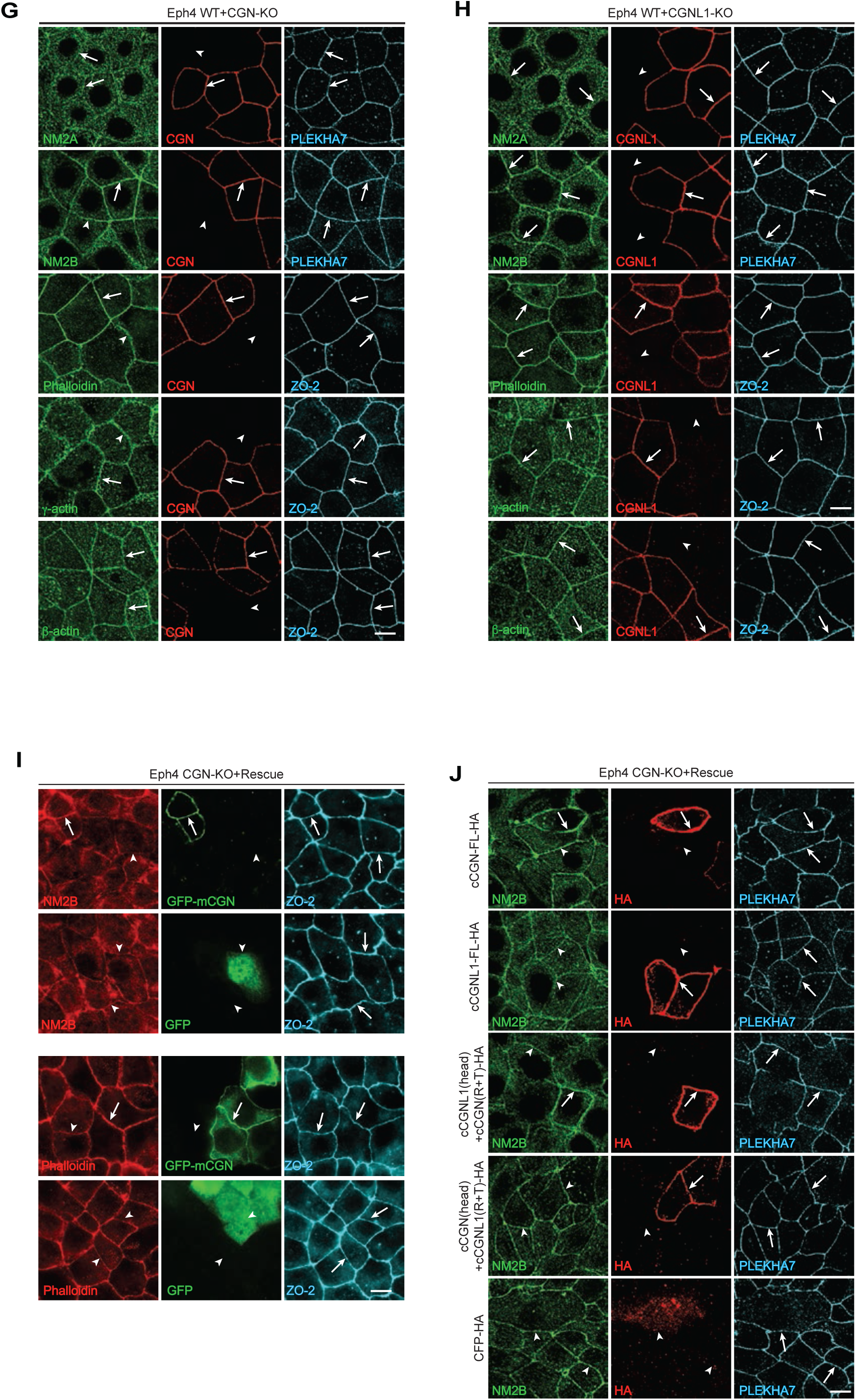
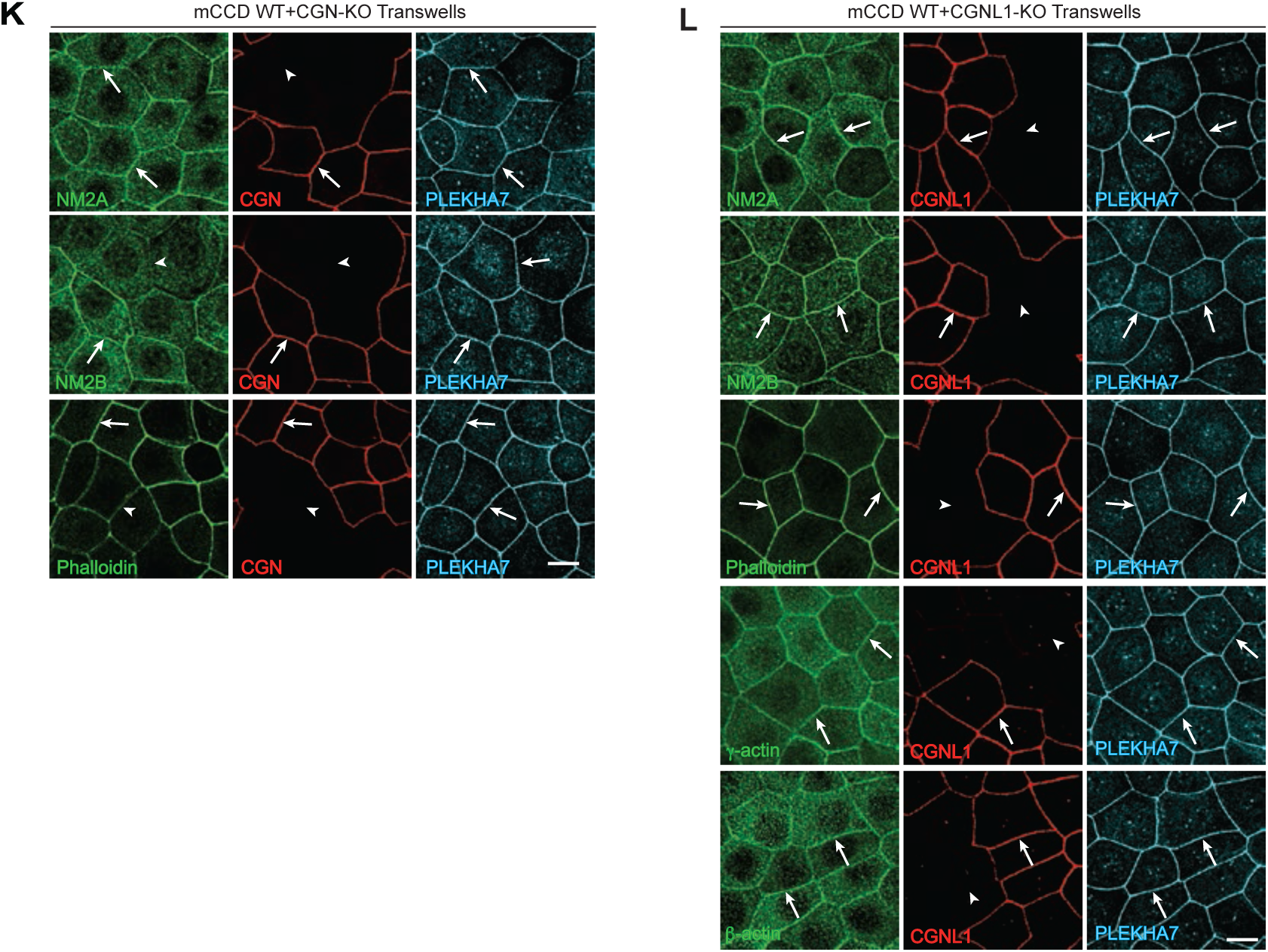
(Related to Figure 2). Cingulin controls membrane tortuosity in MDCK cells and recruits myosin2B (NM2B) to tight junctions of Eph4 and mCCD cells. (A) Quantification of zig-zag index in either CGN-KO or WT MDCK cells rescued with the indicated constructs. (B-C) Immunofluorescent (IF) localization of occludin and the indicated HA-tagged rescue constructs in either WT (B) or CGN-KO (C) MDCK cells. Lines in the occludin channel trace membranes to highlight shape. (D) IF localization of cingulin in MDCK WT cells transfected with the indicated HA-tagged constructs, showing increase in junctional cingulin induced by overexpression of ZO-1. (E) IF localization of NM2A, NM2B, actin filaments (phalloidin) and ARHGEF18 in mixed cultures of WT and CGNL1-KO MDCK cells. NB: CGNL1-KO cells were identified by stable expression of YFP, since endogenous levels of CGNL1 were too low to be reliably detected in WT-KO mixed cultures. Asterisks in (E) indicate diffuse apical (terminal web) labeling (F) IF localization of NM2B, F-actin (phalloidin), and ARHGEF18 in CGN-KO MDCK cells rescued with GFP alone. Either PLEKHA6 or ZO-2 were used as junctional reference marker in MDCK cells, that express low levels of PLEKHA7. (G-H) IF localization of NM2A, NM2B, actin filaments (phalloidin), γ-actin and β-actin in mixed cultures of either WT or CGN-KO Eph4 cells (G) or WT and CGNL1-KO Eph4 cells (H). Either PLEKHA7 or ZO-2 were used as junctional reference marker. (I-J) IF localization of NM2B and actin filaments (phalloidin) in CGN-KO Eph4 cells transfected with GFP-msC-GN-myc and GFP-myc (I) or localization of NM2B after transfection with indicated CGN/CGNL1 full length or chimaeric HA-tagged constructs (J). (K-L) IF localization of NM2A, NM2B, actin filaments (phalloidin) in mixed cultures of either WT or CGN-KO mCCD cells (K) or WT and CGNL1-KO mCCD cells (L). γ-actin and β-actin were examined also in WT-CGNL1-KO mixed cultures of mCCD (L). PLEKHA7 was used as junctional reference marker. Arrows= normal labeling, double arrows=increased labeling, arrowheads=reduced/undetectable/cytoplasmic labeling. Scale bars, 10 μm.

**Figure S3.**
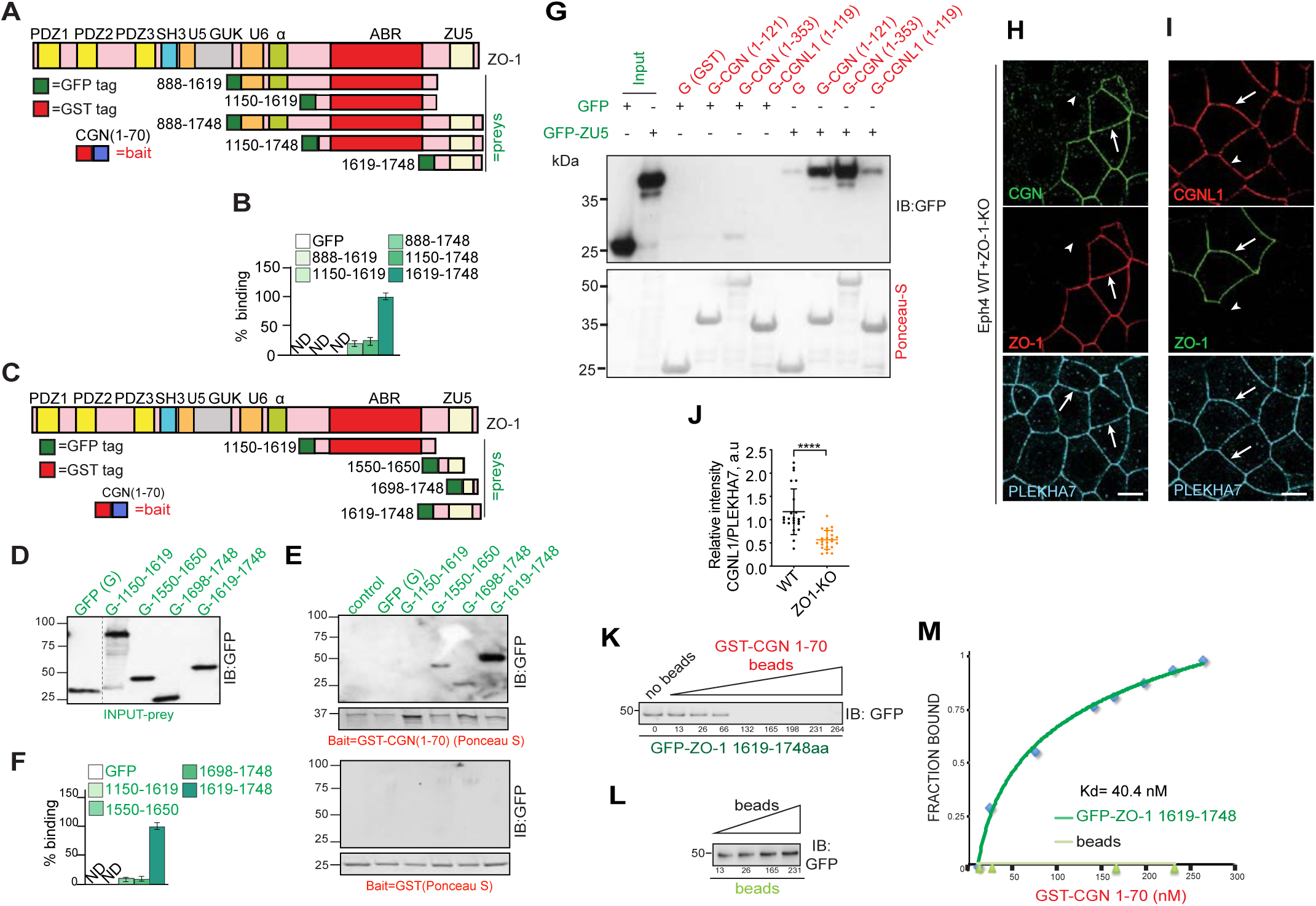
(Related to Figure 3). Binding of cingulin and paracingulin to the ZU5 domain of ZO-1. (A-B) Relative binding of ZO-1 fragments to CGN(1-70). (A) Schematic diagram of ZO-1 with its structural domains: PDZ (Psd-95/Dlg/Z0-1), SH3 (Src-homology3), U5, GUK (guanylate kinase), U6, a, ABR (actin-binding-region), and ZU5. Bait and prey proteins are shown with either red (GST) or green (GFP) tags, respectively. Numbers correspond to the amino acid residues comprised in each construct. (B) Histogram, based on densitometry analysis, showing relative binding of ZO-1 fragments to the CGN(1-70) bait, taking binding of ZU5 (residues 1619-1748) as 100%. (C-F) The integrity of ZU5 domain is required for interaction with cingulin. (C) Schematic diagram of GFP-tagged ZO-1 prey constructs and GST-CGN(1-70) bait used in pulldown. (D) Prey normalization. (E) Immunoblot analysis, using anti-GFP, of pulldowns using either GST-CGN(1-70)(top) or GST (bottom) as baits (PonceauS), and normalized GFP-tagged ZO-1 fragments as preys. (F) Histogram, based on densitometry analysis, showing relative binding of ZO-1 fragments to the CGN(1-70) bait, taking binding of the ZU5 as 100%. (G) Interaction of ZU5 with fragments of cingulin and paracingulin globular head regions. Immunoblot analysis, using anti-GFP, of pulldowns using either GST or the indicated fragment of either CGN or CGNL1 as baits, and GFP-tagged ZU5 as a preys. Numbers correspond to the amino acid residues comprised in each construct. (H-J) Immunofluorescent localization of cingulin (H) and paracingulin (I) in mixed cultures of WT and ZO-KO Eph4 cells. Arrows and arrowheads indicate normal and undetectable junctional labeling, respectively. Scale bars, 10 μm. (J) shows quantification of CGNL1 labeling with respect to the reference junctional marker PLEKHA7. (K-M) Measurement of the affinity of interaction between CGN(1-70) and ZU5. (K) Immunoblot analysis of supernatant depletion assay (STAR Methods). Depletion is achieved by adding increasing amounts of GST-CGN(1-70) beads to the supernatant. Supernatant depletion by beads alone is shown in (L). Numbers below each lane (K-L) indicate concentration (nM) of recombinant proteins. (M) Plots of equilibrium binding isotherms, where fraction of protein bound (total minus remaining) is plotted against the concentration of the GST-CGN(1-70) used for depletion, using either the ZU5 of ZO-1, or beads.

**Figure S4.**
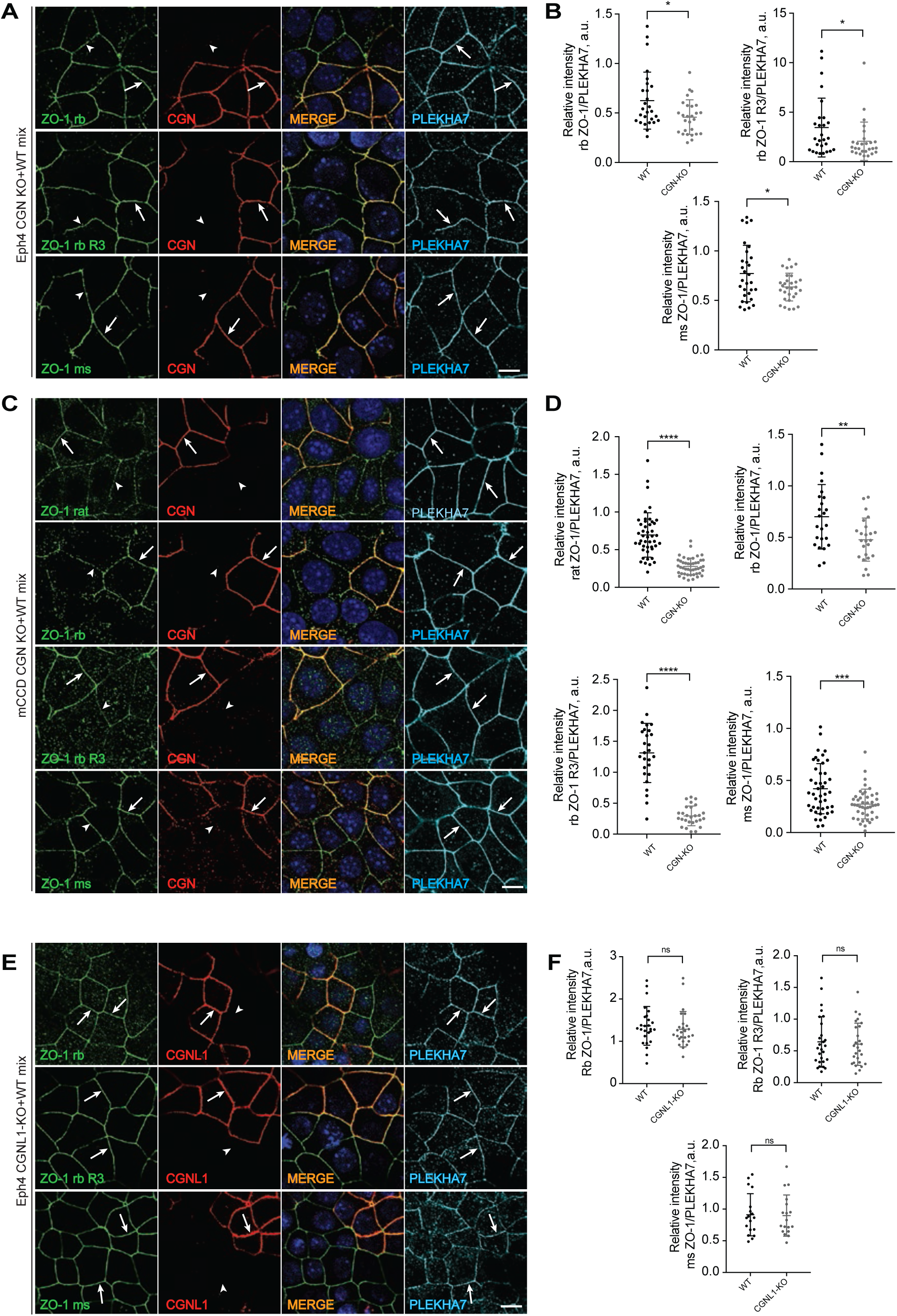
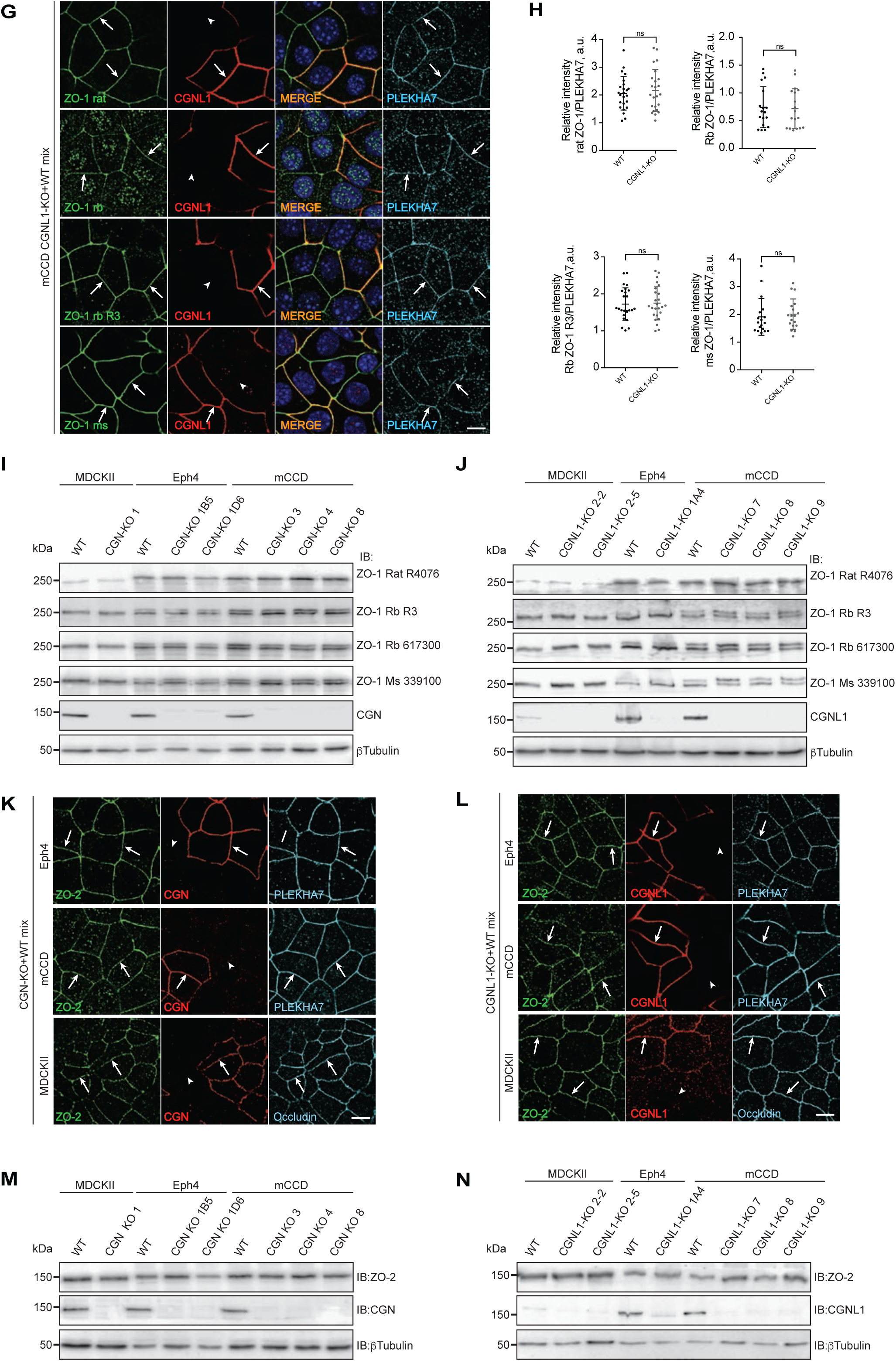
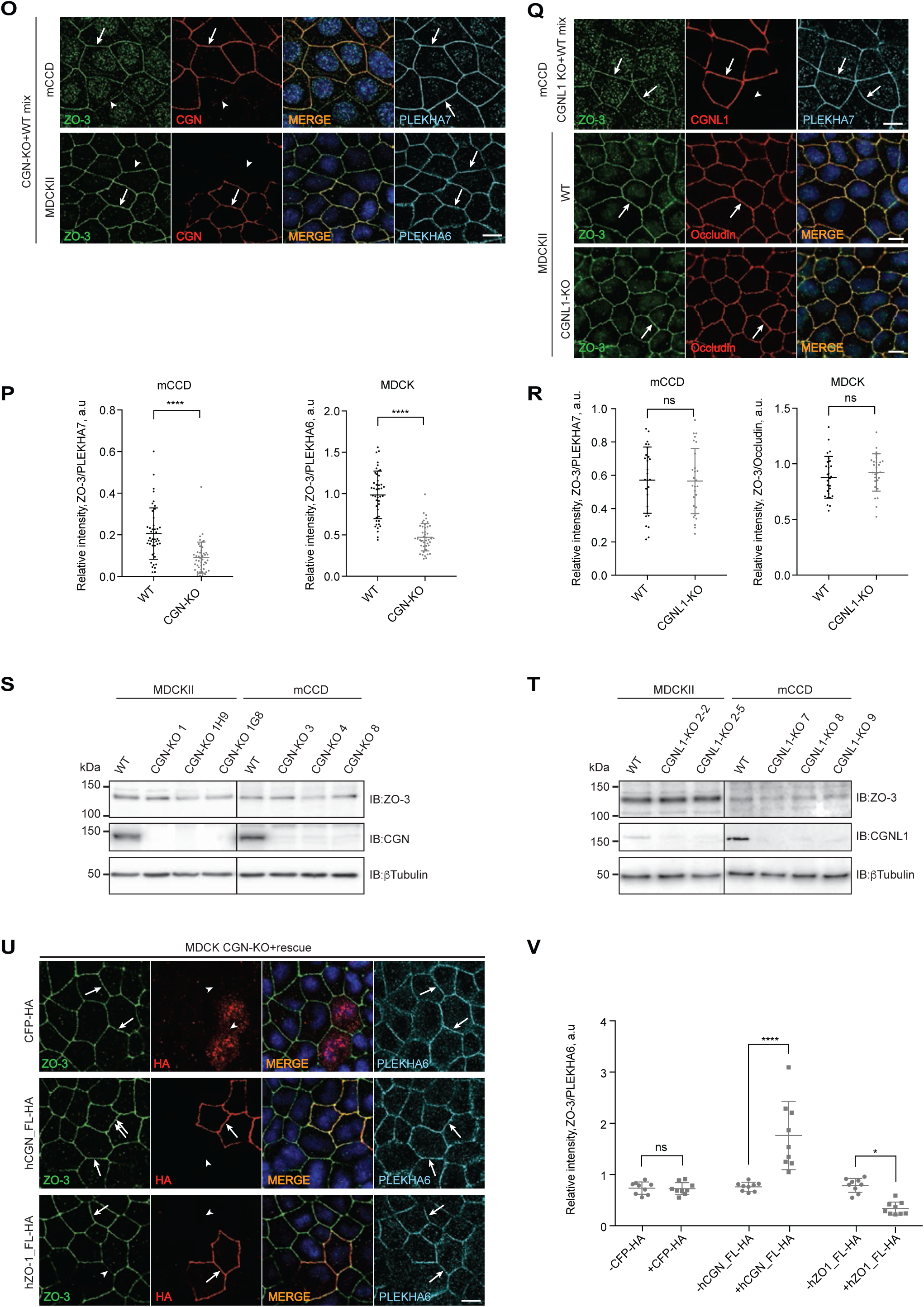
(Related to Figure 4). Cingulin but not paracingulin promote ZO-1 and ZO-3 accumulation at tight junctions. (A-H, K-L, O-R) IF analysis of the junctional accumulation of ZO-1 (A, C, E, G), ZO-2 (K, L), ZO-3 (O,Q) and quantifications of ZO-1 labeling (B, D, F, H), ZO-3 labeling (P, R) with respect to the reference junctional marker PLEKHA7 or occludin (for MDCK) either in mixes of CGN-KO and WT cells (A-D, K, O-P), or in mixes of CGNL1-KO and WT cells (E-H, L, Q-R), using indicated antibodies against ZO-1, ZO-2 or ZO-3. Experiments were done using Eph4, MDCK and mCCD cells, as indicated. Separate cultures of WT and CGNL1-KO MDCK were used to examine ZO-3 labeling (Q-R). (I-J, M-N, S-T). Immunoblotting analysis of lysates of WT and CGN-KO lines (I, M, S), and WT and CGNL1-KO lines (J, N, T) using different antibodies against ZO-1 (I-J), ZO-2 (M-N) or ZO-3 (S-T). Antibodies against either CGN (I, M, S) or CGNL1 (J, N, T), and β-tubulin were used as line phenotype and loading controls, respectively. (U) IF analysis of the junctional accumulation of ZO-3 in MDCK CGN-KO after rescue with either HA-tagged human CGN, ZO-1 or CFP. PLEKHA6 was used as a junctional reference. (V) Quantification of ZO-3 junctional IF signal in MDCK CGN-KO cells after rescue with either CFP-HA or full-length human CGN or ZO-1 constructs (see panels U). Scale bars for IF, 10 μm. Data in plots represented as mean±SD. Signal intensities are shown in arbitrary units (a.u.).

**Figure S5.**
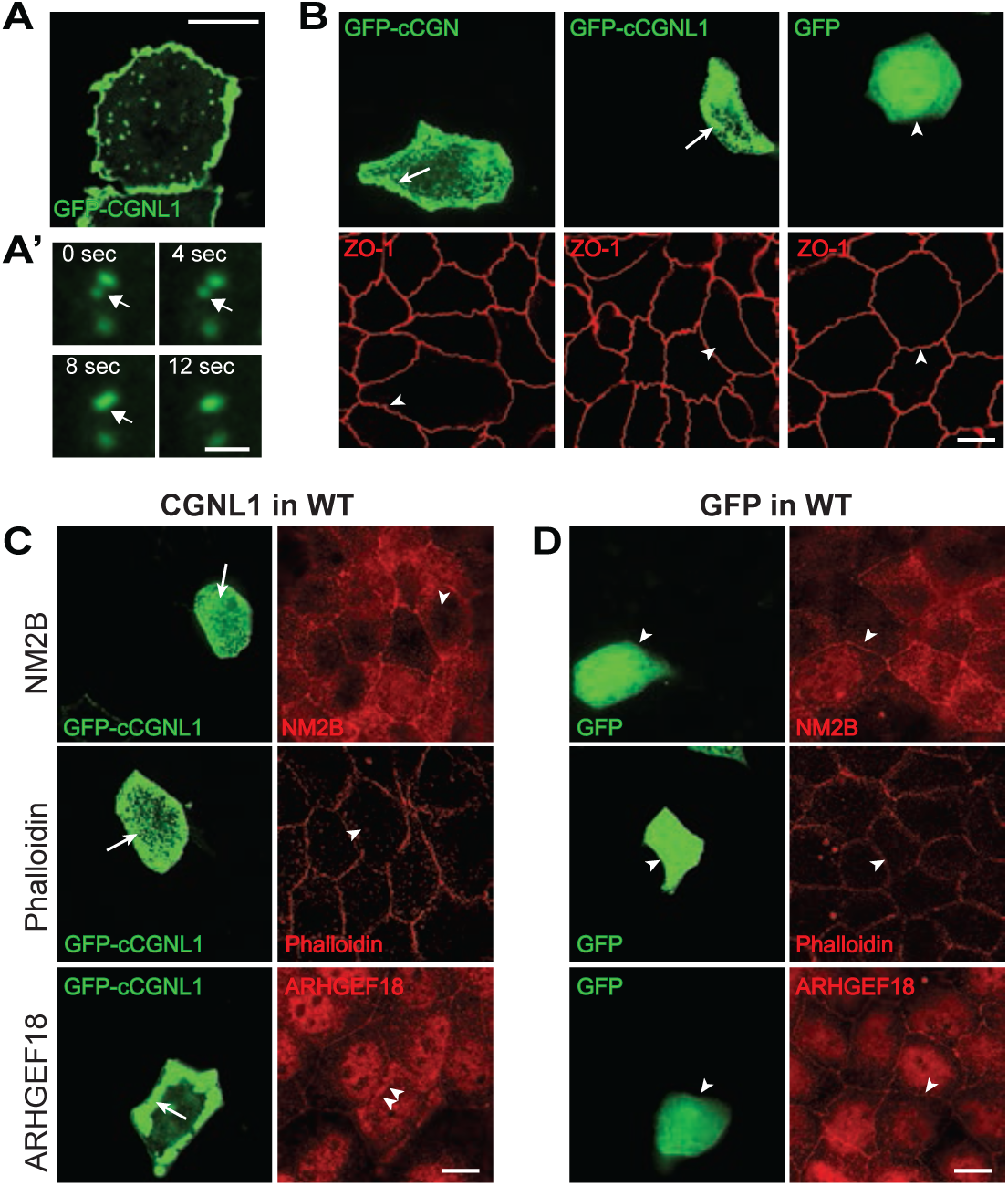
(Related to Figure 5). Cingulin and paracingulin condensates do not recruit ZO-1. (A) CGNL1 form droplets that fuse into larger structures over time (A’), suggesting the formation of phase-separated condensates. (B) IF analysis of ZO-1 in condensates of either GFP-CGN or GFP-CGNL1 or after overexpression of GFP (negative control) in MDCK WT cells. Arrows indicate condensates, arrowheads indicate lack of accumulation of ZO-1. (C-D) IF analysis of NM2B, F-actin (TRITC-phalloidin) and ARHGEF18 in cells overexpressing either GFP-CGNL1 (C) or GFP alone (D). Arrows indicate condensates, arrows and arrowheads indicate co-accumulation or lack of detectable co-accumulation in condensates, and double arrowheads indicate redistributed subcortical labeling, respectively. Scale bars, 10 μm, except for (A) = 0.05 μm.

**Figure S6.**
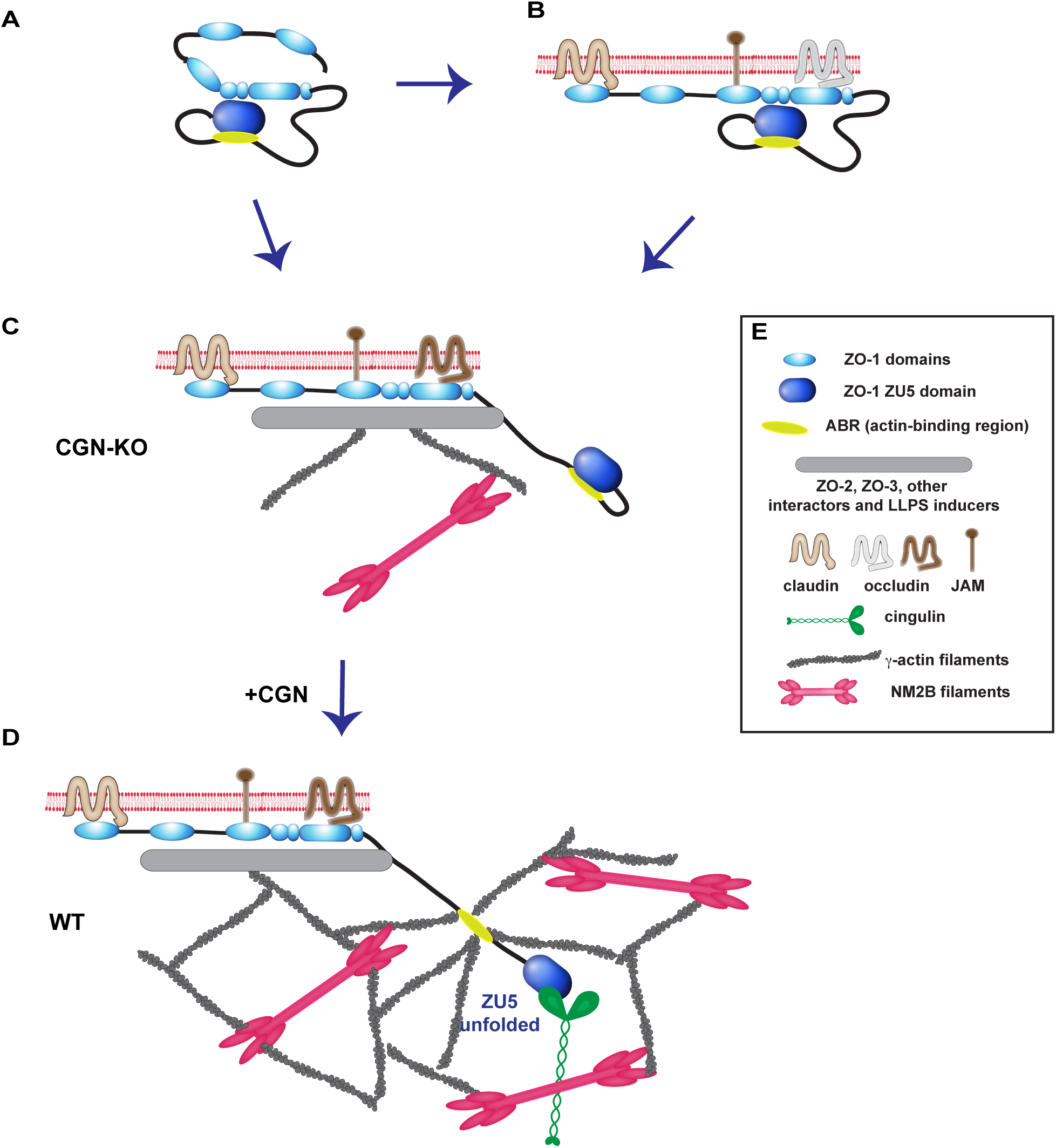
(Related to Figure 2, Figure 3, Figure 5). Scheme linking conformational states of ZO-1 to junction association and interactions with cingulin, actin filaments and other proteins. (A) Cytoplasmic ZO-1 (soluble, folded). ZU5 undergoes intramolecular interactions with ZPSG (Spadaro et al, 2017) and ZU5 domains. (B) Junction-associated ZO-1 in the absence of ZO-2 and cingulin, and with disrupted actomyosin organization/tension interacts with claudins and JAM-A, but does not recruit either occludin or DbpA (Figure 3, (Spadaro et al., 2017)). NB: in the presence of cingulin the ZU5 domain would unfold, but this is not sufficient to induce full ZO-1 stretching. (C) Junction-associated ZO-1 in the presence of interactors of the PDZ2-PDZ3-SH3-GUK and C-terminal domains, some of which may indirectly link ZO-1 to actin and myosin, and additional factors (phosphorylation (Beutel et al, 2019)) that induce liquid-liquid phase separation (LLPS) becomes unfolded and phase separates. However in the absence of cingulin (CGN-KO) the ZU5 domain is folded onto the ABR domain, inhibiting its interaction with actin filaments. (D) The ZO-1 configuration shown in C (CGN-KO) plus the addition of cingulin allows unfolding of the ZU5 domain and organization of the NM2B-γ-actin web. (E) Key for identification of graphical elements.

